# Complementary Structural and Chemical Biology Methods Reveal the Basis for Selective Radioligand Binding to α-Synuclein in MSA Tissue

**DOI:** 10.1101/2025.06.30.662461

**Authors:** Juan C. Sanchez, Ryann M. Perez, Dinahlee Saturnino Guarino, Dhruva Dhavala, Joseph K. Baumgardt, Kyle D. Shaffer, Paul Kotzbauer, Robert H. Mach, E. James Petersson, Elizabeth R. Wright

## Abstract

Fibrillar aggregation of α-synuclein (αSyn) is a hallmark of Parkinson’s disease (PD) and related disorders, including multiple system atrophy (MSA) and dementia with Lewy bodies (DLB). Despite advances in αSyn fibril structural characterization, the relevance of *in vitro* and *ex vivo* structures to patient aggregates remains unclear, particularly for developing therapeutic or diagnostic molecules. Cryo-electron microscopy (cryo-EM) studies of αSyn fibrils with ligands often reveal binding at multiple sites, likely due to high ligand concentrations. Here, various structural and chemical biology techniques were used to characterize αSyn fibrils in the presence of EX-6, a candidate ligand for positron emission tomography (PET) imaging of synucleinopathies. Transmission electron microscopy (TEM) and cryo-EM revealed no significant fibril core changes upon binding. Förster resonance energy transfer (FRET) further demonstrated that the disordered C-terminus was unaltered. Cryo-EM and crosslinking mass spectrometry (XL-MS) identified consistent binding sites, with one (Site 2*) providing a well-defined pocket for high-resolution analysis. Site 2* showed similar residue positioning in MSA patient-derived structures, suggesting MSA selectivity. [^3^H]-EX-6 binding assays demonstrated a 10-fold preference for MSA over PD tissue, with autoradiography further confirming MSA selectivity. Taken together, the combined use of structural and chemical biology techniques provides a comprehensive understanding of EX-6 binding that would not be possible with any single method. Optimization of ligand-protein and ligand-ligand interactions observed in the cryo-EM structure will enable the development of EX-6 as a PET imaging probe for MSA.

## Introduction

The diagnosis of many neurodegenerative diseases is based on a physician’s assessment of the neurological symptoms and impairments present in a patient. Unfortunately, for many neurological disorders, the symptoms exhibited are similar and may lead to misdiagnosis and/or rediagnosis years into treatment. Additionally, several of the clinical symptoms used in diagnosis are not observed until later stages of the disease. This is especially applicable to synucleinopathies, i.e., neurological disorders tied to the overexpression and/or aggregation of *∝*-synuclein (αSyn) in neurons and glia, for which no conclusive diagnostic test is available. These synucleinopathies include Parkinson’s disease (PD), multiple system atrophy (MSA), and Lewy body dementia (LBD), which are all characterized by impaired movement, accompanied by cognitive impairment in LBD and frequently accompanied by autonomic impairment in MSA (1). The lack of diagnostic tools for this subset of neuronal diseases is especially concerning because PD is the second most common neurodegenerative disorder in the United States, behind Alzheimer’s disease (AD). The accuracy of clinical diagnosis of PD ranges from 76% to 92% (2, 3). While there has been recent progress in detecting the presence of aggregated αSyn in biological fluids and skin samples, which may improve diagnostic accuracy for PD, these approaches do not provide information on the distribution of pathology in the brain (4-6). Staining of αSyn aggregates in brain tissue slices has been crucial to understanding the pathology of PD, MSA, and LBD; however, this can only be undertaken postmortem. For example, studies of PD postmortem tissue indicate that the development of dementia in PD is associated with more widespread accumulation of aggregated αSyn, involving limbic and neocortical regions in addition to the subcortical areas such as the substantia nigra (7-9). Thus, there is a need for tools to image the progression of synucleinopathies *in vivo*, which would provide biomarkers for disease progression that can be used to guide the development of disease-modifying therapies. Structural information on the interaction of imaging probes with aggregated αSyn can guide the development of these tools.

αSyn is a 140 amino acid protein that is encoded by the *SNCA* gene and is characterized by three unique domains (10). The first 60 residues comprise the N-terminal amphipathic region, which has been shown to be involved in lipid binding (11). Residues 61-95 are known as the non-amyloid-? component (NAC) domain, which is essential for αSyn aggregation (12-14). Finally, residues 96-140 are considered the C-terminal acidic domain that appears crucial for maintaining the cytosolic form of the protein, and modifications to this region have shown increased propensity for protein aggregation (15, 16). αSyn is predominantly located in the presynaptic terminal and is believed to be essential for regulating presynaptic vesicle fusion and neurotransmitter release (12, 17, 18). In αSyn’s cytosolic and monomeric form, it is entirely disordered and forms a helical structure when bound to vesicle membranes, with some evidence for tetrameric assembly (19, 20). However, through a mechanism that is not fully understood, αSyn aggregates into fibrillar structures that are linked to neurotoxicity (21). Multiple fibrils coalesce to form neuronal inclusions called Lewy bodies and Lewy neurites in PD/LBD, and glial cytoplasmic inclusions (GCIs) in MSA, the histological hallmarks of synucleinopathies (21).

Although there has been incredible progress in determining the molecular details that drive αSyn function and pathology, the question remains: how can one protein be associated with distinct disease states? Recent cryo-electron microscopy (cryo-EM) studies on recombinant αSyn fibrils have shown that several fibril polymorphs can be generated, depending on incubation conditions and the presence of missense mutations or post-translational modifications of the protein (22-26). Histological studies on postmortem brain tissue have shown that the location of αSyn inclusions is tied to different disease phenotypes (21, 27). Structural studies of patient-derived fibrils revealed unique fibril folds, or morphologies, dependent on the disease phenotype, that were not observed for *in vitro* fibril preparations (28-30). Through these structural studies, there is increasing evidence that the clinicopathological heterogeneity of synucleinopathies is linked to differences in αSyn at a molecular level, and that the design of imaging probes for αSyn will need to account for these morphological differences.

Small molecules that specifically bind to αSyn fibrils could serve as positron emission tomography (PET) imaging probes to study disease progression or act as tools for early clinical diagnosis and differentiation of PD from other synucleinopathies, where disease specificity is essential (31). Indeed, such PET probes have proven invaluable for studying AD progression and evaluating the efficacy of therapeutics (32). To develop such PET ligands for αSyn, the Mach and Petersson laboratories employed an *in silico* screening method to identify a series of small molecules that could act as potential αSyn fibril radioligands (33). The development of these compounds began with the identification of several potential binding pockets observed in the solid-state NMR (ssNMR) structure of αSyn fibrils (PDB ID: 2N0A) (34). Several binding sites were hypothesized in this study, shown in **Figure S1** in the Supporting Information (33). Site 2 is located near residues Y39-E46 in the 2N0A structure, and Site 9 is located near residues G86-K96. These sites presented deeper binding pockets than Site 3 (K43-H50), Site 7 (V63-G67), or Site 8 (T81-E83), the only other sites that were surface accessible and in the ordered regions of the fibril. For Sites 2 and 9, a pseudomolecule with ideal binding interactions, termed an Exemplar, was generated, and an ultra-high-throughput *in silico* screen was performed to identify commercially available molecules that best matched the Exemplar. An initial set of 17 compounds was purchased and used in competitive binding assays to assess their ability to displace compound Tg-190b or BF2826, molecules determined to selectively bind to Sites 2 and 9, respectively, through photo-crosslinking mass spectrometry (XL-MS) experiments (33). The results showed that compound 6, termed Exemplar-6 (EX-6, **Figure 1C**), has a binding affinity of 9 nM against Site 2 and could serve as a potential radioligand for imaging synucleinopathies (33).

**Figure 1.**
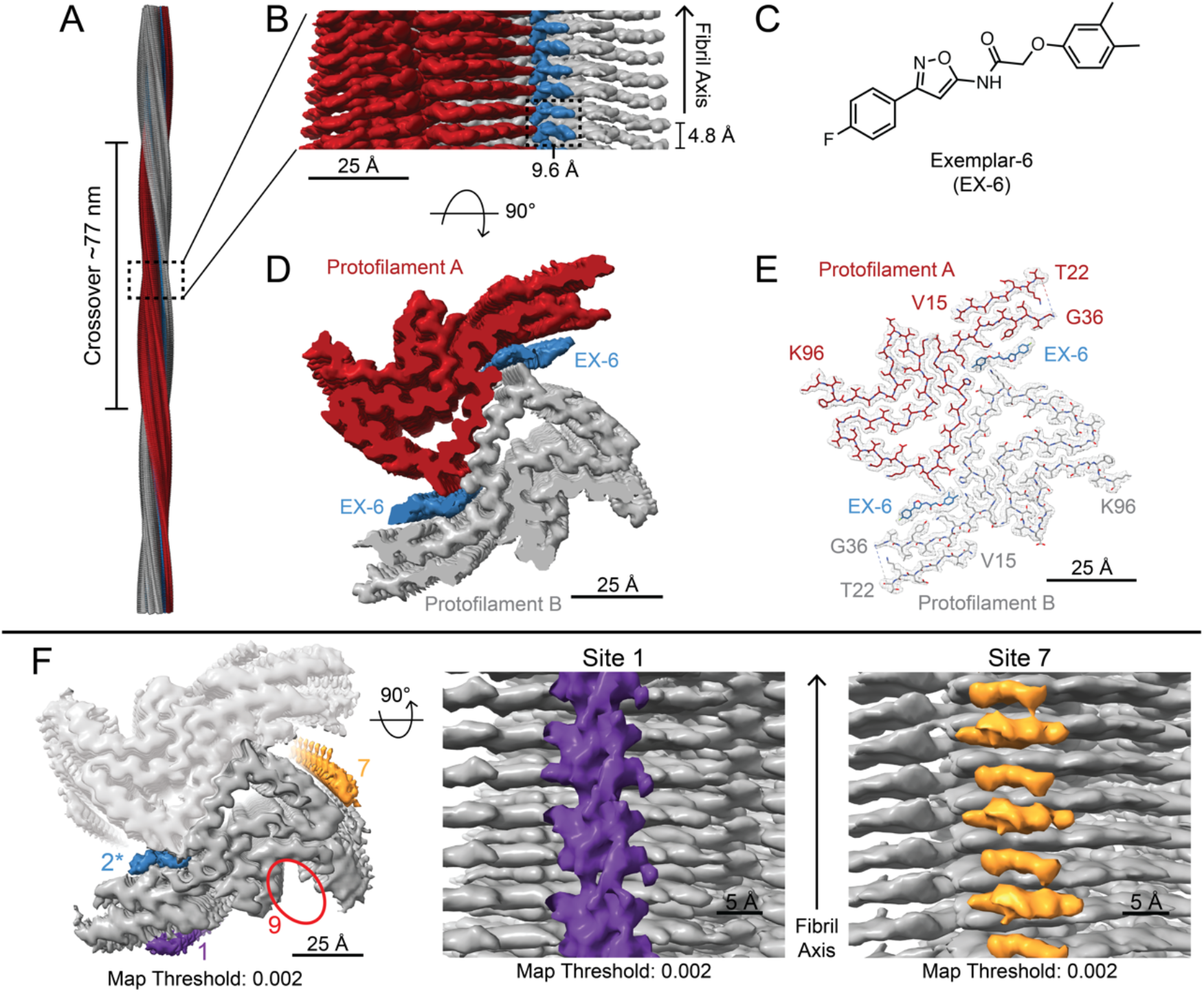
Cryo-EM Data for EX-6 Bound Fibrils A. Multiple cryo-EM maps aligned and stacked together along the fibril axis for easier visualization of the twisting nature of the fibrils. B. A close-up of the cryo-EM map shows the primary ligand binding pocket (Site 2*). The map is segmented for distinction between protofilaments and the ligand. The map threshold is set to 0.003. C. Diagram of Exemplar-6 (EX-6). D. Map from B rotated 90° to show a cross-section of the fibrils, the map threshold is set to 0.003. E. Map and model overlay of the cross-section view showing good agreement between the map and model; only one fibril layer is shown. F. Cross-sectional view of fibrils with map threshold set to 0.002, showing additional density attributable to lower-affinity ligand binding sites, denoted Site 1 and Site 7. No observable density at Site 9 (red circle).

Although the *in silico* methods proved helpful in identifying lead compounds, the input model was that of a single protofilament. The latest structural data show that *in vitro* αSyn fibrils commonly exist as multi-protofilament structures, and this can result in a rearrangement of Site 2 and Site 3 (backbone rotation that places Y39 and H50 on the same face of the protofilament) to form a site that we will refer to as Site 2*. Moreover, the similarities and differences between these structures and those obtained from *ex vivo* preparations of fibrils from PD or MSA patients raise essential questions about how data for *in vitro* fibrils can usefully inform the development of molecules targeting αSyn aggregates in patients. Thus, our goal was to use cryo-EM to determine a high-resolution structure of EX-6 in complex with αSyn fibrils, and complement these data with a suite of additional structural, computational, chemical biology, and radiology methods to provide a full understanding of ligand binding.

Our complementary, yet orthogonal, research methods yielded multiple lines of evidence indicating that EX-6 primarily binds at Site 2* in a well-defined orientation, providing a clear explanation for compound selectivity in tissue. Cryo-EM data provided consistent models for the placement of the ligand in Site 2*, and the cryo-EM data could be refined to generate a high-resolution model, revealing key ligand-protein and ligand-ligand interactions. Cryo-EM also showed that the fibril core was unaltered upon ligand binding, and Förster resonance energy transfer (FRET) showed that the disordered regions that might regulate binding site access were also unaltered. XL-MS data corroborated the cryo-EM findings, including the presence of weaker alternate binding sites, and showed that the binding sites are maintained down to nM concentrations. Comparison of the Site 2* binding pocket to the same region in *ex vivo* structures of αSyn fibrils from PD/DLB or MSA patients provides a clear structural rationale for the MSA selectivity of EX-6 observed in autoradiography studies with patient tissue samples. The comprehensive picture of ligand binding, assembled from an array of methods, can be used to guide the development of new EX-6 derivatives as MSA PET imaging probes and serves as a paradigm for investigating other ligands in synucleinopathies and related amyloid diseases.

## Results and Discussion

### Fibril sample preparation

Since we wished to employ a multimodal approach to enable a comprehensive understanding of ligand binding, it was essential to ensure that fibril morphology was consistent across experiments, as fibril morphology can be variable even among seemingly identical preparations. Therefore, the same fibril samples were shared among laboratories and used in multiple types of experiments to cross-validate findings. Thus, ligand binding information should be directly comparable across these studies. While FRET studies were conducted using a different fibril preparation due to the incorporation of a fluorescently labeled protein, XL-MS studies conducted on these fibrils confirmed that the binding sites were maintained. The observed polymorph is also consistent with literature reports from other laboratories using similar conditions (35).

### Fibril structure is unaltered upon ligand binding

An analysis of NS-TEM imaging data confirmed that the addition of the EX-6 ligand to the fibrils did not result in major structural changes to the fibrils themselves (**Figure S2**) The fibril widths were similar for both apo and EX-6 bound fibrils, measuring 11.0 nm and 10.6 nm, respectively (**Figure S2**), measured from the cryo-EM data (36). Furthermore, there were no obvious indications in NS-TEM images that EX-6 caused fibrils to dissociate or break down, consistent with previous studies (33, 37). However, due to the molar excess of ligands over fibrils, we observed ligand aggregates on the negative-stain grids that were not present in the apo grids (**Figure S2**, red circles). We suspect these are due to the excess ligands present in the sample adsorbing the stain and adhering to the carbon film of the grid. Further support for the lack of fibril perturbation upon ligand binding came from FRET and cryo-EM studies, described below. Together, these results suggest that at a higher level, the ligands do not alter the fibril structure in any significant way, as expected based on previous fibril analysis using fluorescence polarization (33, 37).

### Cryo-EM structural characterization of fibrils

The cryo-EM data were consistent with NS-TEM images and showed several twisting fibrils with a few overlapping regions per micrograph, which we incorporated into the data processing. After preprocessing steps, 2D classification for both the apo and EX-6 ligand data revealed one primary fibril form with an estimated crossover distance of 77 nm (**Figure 1A**). Cryo-EM data processing, described in detail in the Supporting Information, resulted in a final reconstruction to a global resolution of ∼2.0 Å for the apo structure (**Figure S3**, EMD-45639) (36) and ∼2.6 Å for the ligand-bound structure (**Figures S4-S5, Table S1**, EMD-70093). A model comprising six subunits per protofilament was built into the central region of each map. Of the 140 amino acids in the αSyn protein, 71 residues were unambiguously modeled with excellent side chain density for the apo structure, and 70 residues for the ligand-bound structure. In our model, an amyloid “Greek Key” structural motif is present, as observed in previous αSyn models (35), and is comprised of residues G36 to K97 for the apo structure, and residues G36 to K97 for the ligand-bound structure. In addition, our well-refined ligand-bound EM map contained an island of density near the N-terminal region where residues V15 to T22 fold back towards the fibril core as in the apo structure (**Figure 1E, Figure S3**) (36). The intra-protofilament rise is ∼4.8 Å, and the two protofilaments interact to form a stable fibril core with an additional pseudo-2_1_ screw symmetry present between the two protofilaments, as observed in the apo map (**Figure 1B, Figure S3**). A further well-refined density attributed to the EX-6 ligand at Site 2* allowed for docking of 12 EX-6 molecules along the fibril axis, 2 per fibril layer (**Figure 1B**). Thus, the refined cryo-EM map reveals two twisting protofilaments that create two Site 2* pockets per fibril layer where EX-6 ligands bind in a 1:1 fashion (**Figure 1B, 1D)**.

### Protein interactions stabilizing the observed fibril polymorph

Our molecular models align with previously solved recombinant αSyn structures comprising two protofilaments that interact and form an interface including residues H50 and E57 between opposing protofilaments, also termed the preNAC interface (**Figure 1E, Figure S3)** (22, 23). Salt bridges at the ends of the preNAC interface stabilize the fibril core, specifically between H50 and E57 from opposing protofilaments. Hydrophobic interactions between opposing valine and alanine residues at this interface further stabilize the fibril core, resulting in a steric zipper that runs the length of the fibril axis **(Figure 1E, Figure S3)**. The average buried surface area at the preNAC interface for both the apo and the EX-6 ligand-bound fibrils is ∼176 Å^2^, with a solvation free energy of -79 kcal/mol between fibril layers (38-40). These results are consistent with our high-level observations that ligand binding does not alter the fibrils and that binding occurs at solvent-accessible regions.

For several published models, the N-terminus appears to be in slightly different conformations (22, 23, 35). In our case, a well-resolved island of density allowed us to model residues V15 to T22, showing that the N-terminal region folds back towards the fibril core (**Figure1, Figure S3)**. This region is stabilized by hydrogen bonds between the V16 backbone carbonyl oxygen and the S42 side chain. Additionally, interdigitation of A17, A19, L38, and V40 forms a hydrophobic interface that further stabilizes this region.

The organization and stabilization of the N-terminus results in a conformation with a cationic, solvent accessible pocket that we will refer to as Site 2* for its similarity to the merged Site 2/Site 3 binding site noted in previously reported two protofilament αSyn fibril polymorphs. Site 2* is created by residues G36 to H50 located on one protofilament and E57 to K60 from the second protofilament (**Figure 1F**). An identical, symmetry-related cavity forms on the opposite side of the protofilament interface (**Figure 1**). Interestingly, this cavity contains three lysine residues (K43, K45, and K58) that are solvent-exposed and are likely stabilized in the apo structure by phosphate ions in the fibrilization buffer. These interactions may also explain why the N-terminus is well resolved in our structure, as this portion of the protein is well-stabilized and buttresses the V15 to T22 segment. These three positively charged lysine residues, together with H50 and K60, create an observed cationic pocket at Site 2* that will be explored in further detail.

### Cationic pocket serves as the primary binding site for EX-6

We observed an additional density at the solvent-accessible cationic Site 2* pocket in the EX-6-bound cryo-EM map that was not present in the apo map (**Figure 1**) (36). Initial docking of the EX-6 ligand showed a good fit between the ligand model and the map. The ligand is positioned perpendicular to the fibril axis, with the dimethyl-benzene ring pointed towards the fibril core and the fluorinated ring pointed towards the solvent region (**Figure 2**). Along the entirety of the map, ligands are stacked in this cationic pocket (**Figure S6**), creating a series of ligand-protein and ligand-ligand interactions that provide a molecular rationale for the high binding affinity of EX-6 to these αSyn fibrils (**Figure 2**).

**Figure 2.**
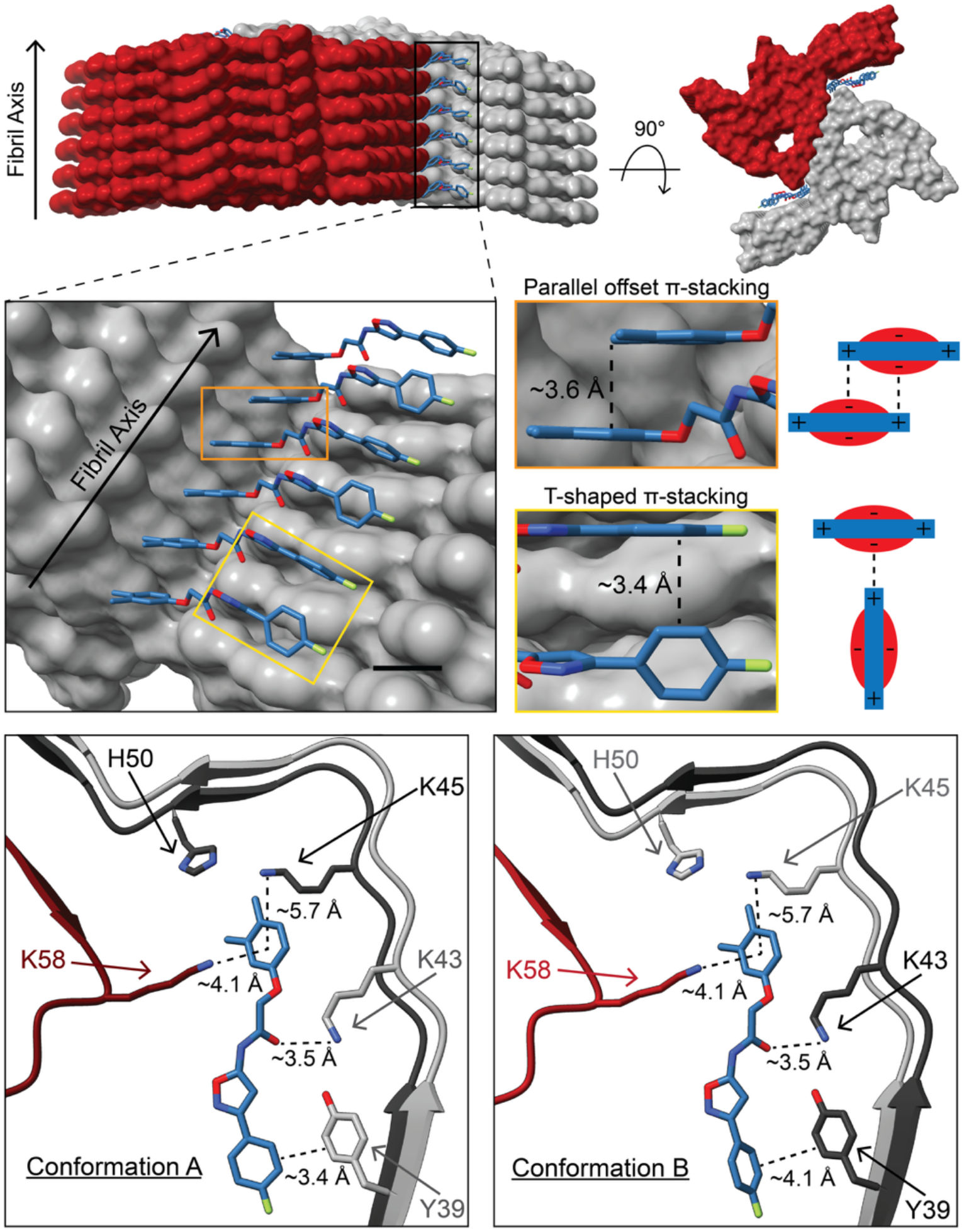
EX-6 Binds Through a Network of Ligand-Protein and Ligand-Ligand Interactions. Top: Site 2* pocket binds EX-6 ligands in two conformations. Middle: The EX-6 dimethyl ring is angled, allowing for ligand-ligand edge-to-edge π-stacking interactions, and alternating conformations of the EX-6 fluorinated ring allow for ligand-ligand T-shaped π-stacking. Middle Right: Illustrations of edge-to-face and edge-to-edge interactions of aromatic rings highlighting alignment of electron-poor aromatic CH and methyl groups with electron-rich π-systems. Bottom: The EX-6 dimethyl ring is further stabilized by ligand-protein cation-π interactions with lysine-45 and lysine-58. Tyrosine-39 and the EX-6 fluorinated ring form a network of π-stacking interactions to stabilize the solvent-exposed region of the ligand.

Comparisons between the apo model and the EX-6 ligand model reveal that side chains for K43 and K45 are slightly displaced to allow for binding of the EX-6 ligand. Upon ligand binding, K43 is rotated towards Y39, forming a hydrogen bond between the side chains that is not observed in the apo structure. K45 is rotated slightly towards the fibril core, creating sufficient space for the dimethyl-benzene ring of the EX-6 ligand in this pocket. Although overall the ligands are positioned perpendicular to the fibril axis and are in a similar plane with the αSyn ?-strands, the dimethyl-benzene ring is rotated ∼60° from this plane (**Figure 2**, orange box). This rotation allows for cation-π interactions between K45 and the dimethyl-benzene ring, measuring ∼5.7 Å in length (**Figure 2**). On the opposite face of the dimethyl-benzene ring, there is a strong cation-π interaction with K58 measuring ∼4.1 Å in length (**Figure 2**). The ligand is likely displacing a negatively charged phosphate in the fibril core, and this is hypothesized to be energetically favorable due to the electron-rich dimethyl-benzene ring providing an additional cation-π interaction (**Figure 2**). In addition to these protein-ligand interactions, the stacked dimethyl-benzene rings form parallel-displaced, or offset, π-stacking interactions measuring ∼3.6 Å from ring edge to ring edge, a geometry that is favorable for electrostatic interactions between the electron rich centers of the rings and the electron poor aromatic hydrogens (**Figure 2,** orange box). These stabilizing interactions at the dimethyl-benzene ring provide a mechanistic explanation for the high binding affinity properties observed in EX-6. Our cryo-EM processing reveals that the dimethyl-benzene ring is positioned in a single conformation throughout the fibril.

### Alternating ligand conformations stabilize EX-6 along the fibril axis

While the dimethyl-benzene ring remained static, the fluorinated benzene ring displayed alternating conformations. In one conformation, the fluorinated ring is in a plane with the αSyn ?-strand at the corresponding fibril layer, then in the adjacent layer, the fluorinated ring of the ligand is rotated by ∼110° (**Figure 2,** yellow box). This pattern is repeated every two fibril layers. The alternating rotation of these rings enables t-stacking interactions to occur between neighboring fluorinated rings along the fibril axis, thereby aligning electron-poor aromatic hydrogens above the electron-rich aromatic rings (**Figure 2,** yellow box). On average, these t-stacking interactions are ∼3.4 Å in length and appear crucial to stabilizing the solvent-facing portion of the ligand. A similar t-stacking interaction is also observed between Y39 and the fluorinated ring that is in plane with the ?-stands measuring ∼3.4 Å in length (**Figure 2**). Offset π-stacking interactions between Y39 and the rotated fluorinated ring (perpendicular to the ?-stands) are also present, with the edges of the rings ∼4.1 Å apart (**Figure 2**). Upon EX-6 binding, K43 rotates and forms a hydrogen bond to Y39, further stabilizing the atoms in this region. The alternating conformations of the fluorinated ring appear to be crucial in holding the ligand in the Site 2* pocket. Together, the ligand-ligand and ligand-protein interactions at the dimethyl-benzene ring and the fluorinated ring form a network that stabilizes the EX-6 ligand at this primary binding site.

### FRET with EX-6 bound

While our structural methods can report on ligand binding directly to the fibril core, they cannot easily discern the influence of these ligands on the disordered regions. Therefore, we employed FRET analysis by strategically placing fluorophores in the disordered regions to report on the global conformations of the fibril after molecule binding. Fluorophores were placed at donor-acceptor positions 94-9, 114-9, and 125-9, selected as sites that would not perturb ligand binding based on the cryo-EM data, with the donor being acridonylalanine (Acd) incorporated using unnatural amino acid mutagenesis (41) and the acceptor being boron-dipyrromethane (BODIPY) attached to a Cys residue via a maleimide linkage (37). The Acd/BODIPY αSyn, Acd-only control αSyn, or BODIPY-only control αSyn constructs were incorporated as 5% of the total monomer during aggregation so that FRET changes would not be influenced by interactions with neighbors (37, 42). FRET measurements indicate that the interchromophore distances remain unchanged after 4 or 24 hours of incubation with EX-6 (**Figure S7-S8, Table S2-S3**). Therefore, it is unlikely that the disordered regions are affected, which is important as they can influence binding site accessibility and protofilament interactions.

### XL-MS of an EX-6 diazirine derivative

Since the cryo-EM studies were performed at super-stoichiometric ligand concentrations, we sought to use XL-MS as a complementary technique to identify EX-6 binding sites at both saturating concentrations and concentrations closer to the low nM range used in PET imaging and radioligand binding studies. Diazirines are used in photo-crosslinking experiments to identify interactions of biological binding partners, including residue-level information gained through MS2 sequencing of crosslinked peptide fragments. Diazirines can be excited with 365 nm light, whereupon they will rearrange to form a diazo intermediate, which can insert into protein side chains, particularly of Glu and Asp residues, or extrude dinitrogen to form a carbene, which can react with a broader set of amino acids (**Figure S9**) (43, 44). A diazirine containing EX-6 analog (EX-6-CLX) was synthesized to confirm Site 2* binding and to investigate the binding sites associated with the other densities observed in the EX-6 cryo-EM data (**Figure 1F)**. After incubation with 500 nM EX-6-CLX and irradiation, the fibrils were disaggregated and subjected to trypsin treatment. Trypsin digests revealed the E46-K58 as the primary crosslinked peptide signal, followed by E11-K22 (**Figure 3A, 3C**). These peptide fragments correspond to Site 2* and Site 1, respectively. MS2 fragmentation of the Site 2* tryptic peptide revealed primary crosslinks to E57 (**Figure 3B**) and lower intensity peaks from crosslinking to H50 (**Figure S16**). While insertion at E57 is certainly influenced by the reactivity of carbenes with acidic residues, it is also consistent with the expected positioning of the crosslinking group based on the orientation of the dimethylbenzene ring in the cryo-EM structure (**Figure 3D**). MS2 data for other peptides are shown in the Supporting Information (**Figure S13-S15**). At higher (100 µM) ligand concentrations, additional crosslinks were observed at Q24-K32 and E35-K43 (**Figure S9**), again consistent with binding to Site 2* and Site 1 (see **Tables S6-S7** for complete MS peak listings). We also performed crosslinking with αSyn monomer. At 100 µM concentrations, some low intensity crosslinking was observed at similar sites to those observed in fibrils (**Figure S10**). However, at 500 nM concentrations, no crosslinking was observed (**Figure S11**). Therefore, it is likely that the crosslinks we observe for fibrils are specific and due to ligand-protein interactions. While the XL-MS results support the idea that Site 2* is the primary binding site, they also strongly identify other sites that are consistent with the additional cryo-EM density at Site 1 (**Figure 3D**). Therefore, we investigated those sites further to attempt to place ligands in these pockets of electron density.

**Figure 3.**
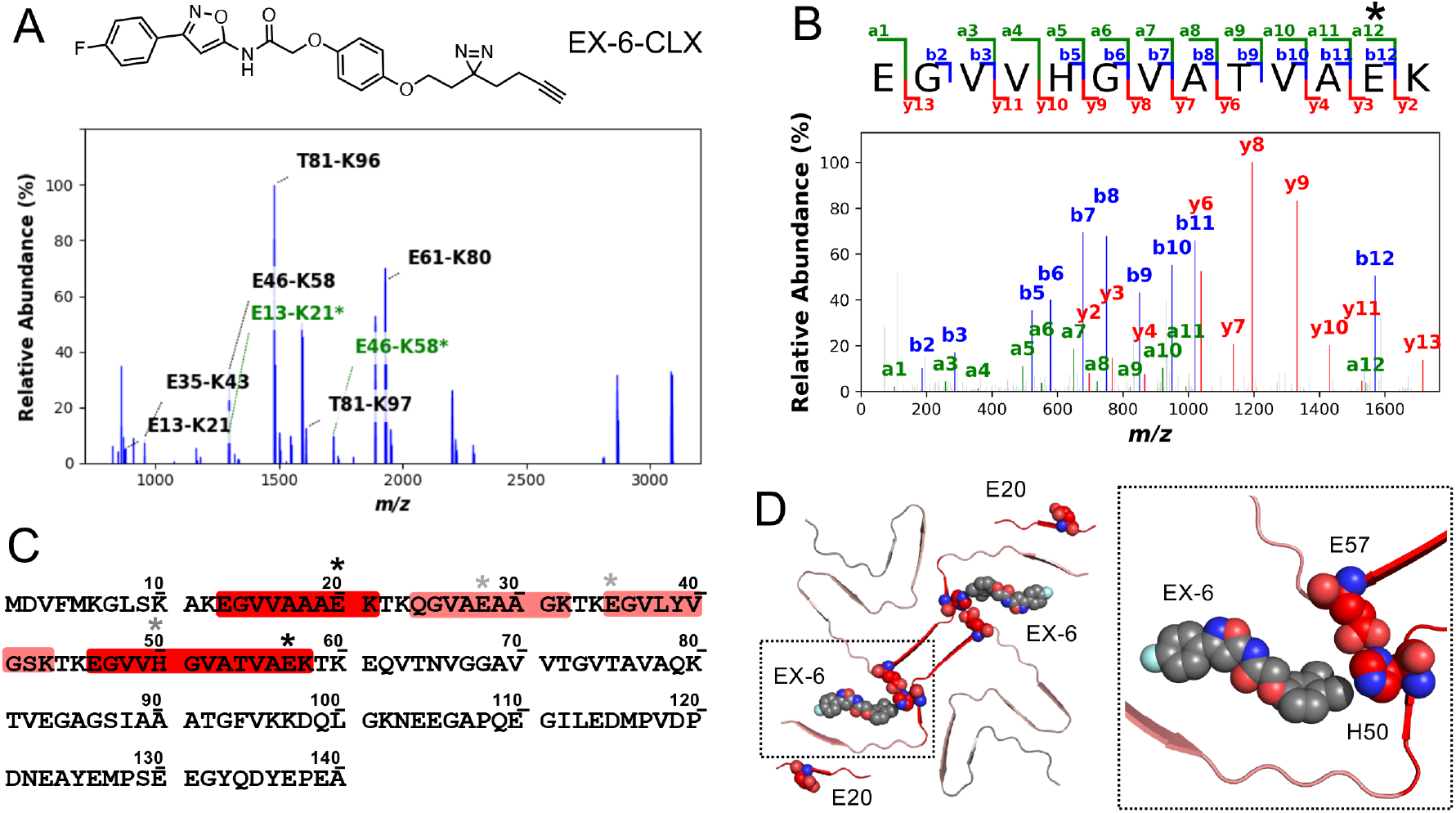
XL-MS of EX-6-CLX A) Trypsin digest MS spectra with crosslinked peaks highlighted in green B) MS2 spectra for the tryptic Site 2* peptide E46-K58, identifying E57 crosslink site (indicated by *) C) Crosslinked peptides mapped on to αSyn sequence with shading intensity indicating identification at 500 nM EX-6 (red) or 100 µM only (pink) (* indicates MS2-identified crosslink site) D) Heat map showing identified peptide fragments (colored as in the sequence diagram) and crosslink sites (spheres) on EX-6 cryo-EM structure.

### Analysis of alternate binding sites in cryo-EM

At lower volume thresholds, there are two additional sites per protofilament that may be additional EX-6 ligand binding sites. Unfortunately, the cryo-EM density at these regions is not resolved well enough to dock and refine the EX-6 molecule unambiguously. The better resolved primary binding site, Site 2*, is present in all EX-6 maps, and the Site 2* ligand density was further improved by additional optimization of helical parameters. These results suggest that the primary binding site is highly occupied across the particle set, and binding occurs in a specific manner at this site. For the potential secondary binding sites, we opted not to refine a molecule into the densities present at Sites 1 and 7, due to the lower local resolution at these sites. However, we can deduce that at Site 1, the ligand appears to be located parallel to the fibril axis (**Figure 1F)**. The ligand at this site likely spans four helical layers and may interact with E20, consistent with XL-MS identification of a crosslink at that residue. At Site 7, the ligand density appears perpendicular to the fibril axis (**Figure 1F**), in-plane with the protein backbone, near Q62 and T64. Due to the low resolution of the map, we were unable to determine the orientation of the ligand at this site. Consistent with the weakness of this interaction, no clear evidence of crosslinking at Site 7 was observed, even at 100 µM EX-6-CLX concentrations. Site 9 does not contain density that could be attributed to a ligand (**Figure 1F**). Thus, XL-MS and cryo-EM are consistent in identifying Site 2* as a high affinity and specificity binding site for EX-6.

### Radioligand binding

To understand the relevance of our EX-6 structural studies to binding αSyn fibril polymorphs populated *in vivo*, we employed radioligand studies in patient brain tissue slices and homogenates. First, we synthesized a tritiated version of EX-6 ([^3^H]EX-6) from a brominated precursor and performed saturation binding assays in brain homogenates from MSA and PD patients with high αSyn pathology scores. As one can see in **Figure 4A**, [^3^H]EX-6 binds in MSA homogenates with a 10-fold higher affinity than in PD homogenates. This is consistent with the relative similarity of the Site 2* region in our structure and in *ex vivo* fibril structures from MSA patients. While several MSA fibril folds have been observed (PDB IDs 6XYO, 6XYP, 6XYQ) with different folds within each protofilament and different protofilament-protofilament interfaces, the Site 2* region adopts a consistent fold that has an RMSD of 1.1-1.3 Å over the G41 to H50 region (**Figure 5**). In contrast, the same region exhibits a 5.1-5.2 Å RMSD when compared to the *ex vivo* PD/DLB fibril cryo-EM structure or the corresponding region in the ssNMR structure of fibrils templated with PD patient αSyn (**Figure 5**). While some questions remain regarding the role of the H50 to E57 region at the protofilament interface in our structure, which is not present in the MSA patient fibril structures, these comparisons provide a structural rationale for the observed radioligand binding selectivity.

**Figure 4.**
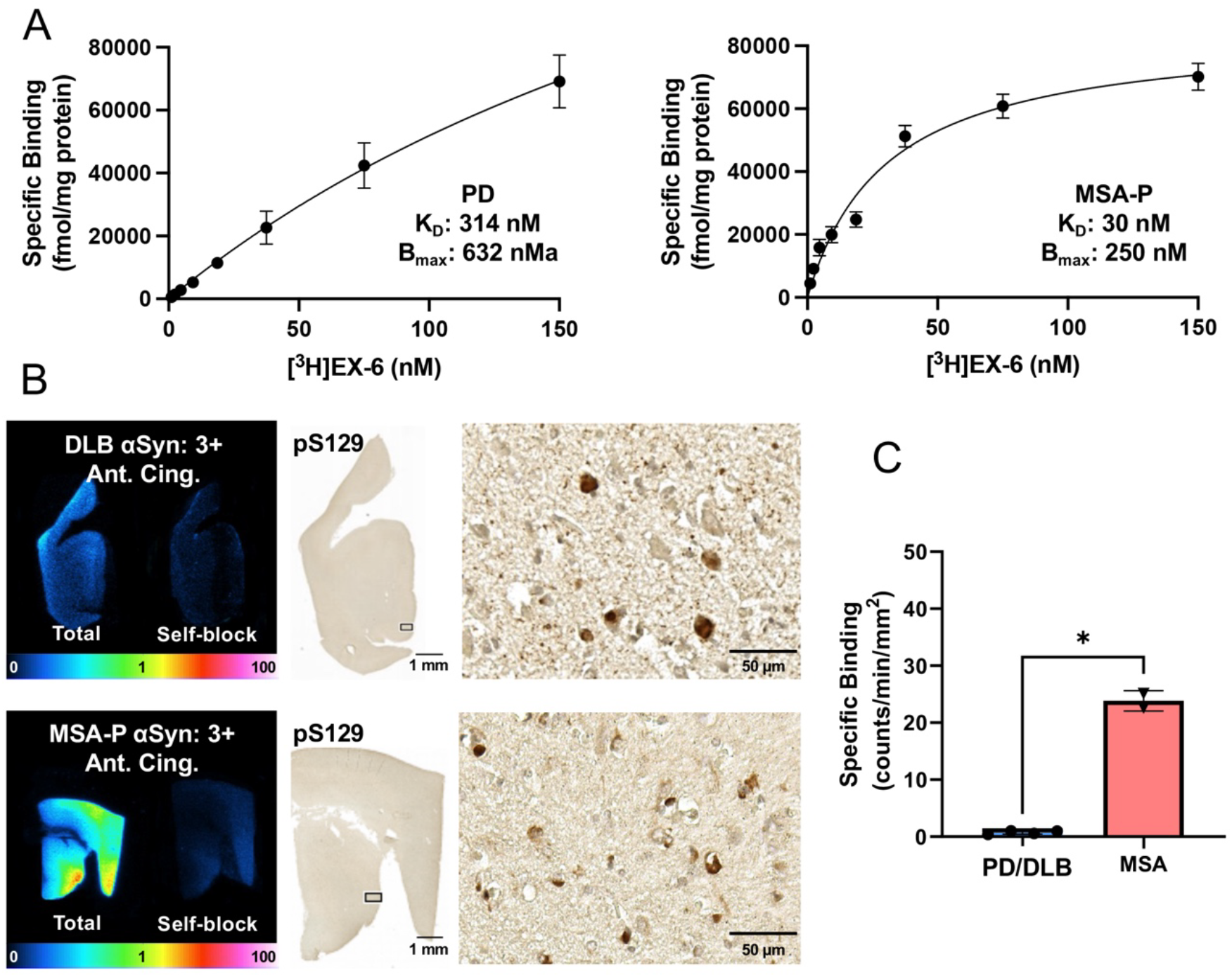
Radioligand Binding Data. A) Dissociation constant (K_D_) for [^3^H]EX-6 determined by saturation binding in PD and MSA patient tissue homogenates, B) Tissue autoradiography studies with [^3^H]EX-6 demonstrating selectivity for binding to MSA samples (MSA-P shown, MSA-C in Supporting Information) and DLB (additional DLB and PD cases in Supporting Information), C) Specific binding in autoradiography experiment shows a >10-fold selectivity for binding to MSA tissues over binding to PD or DLB tissues.

**Figure 5.**
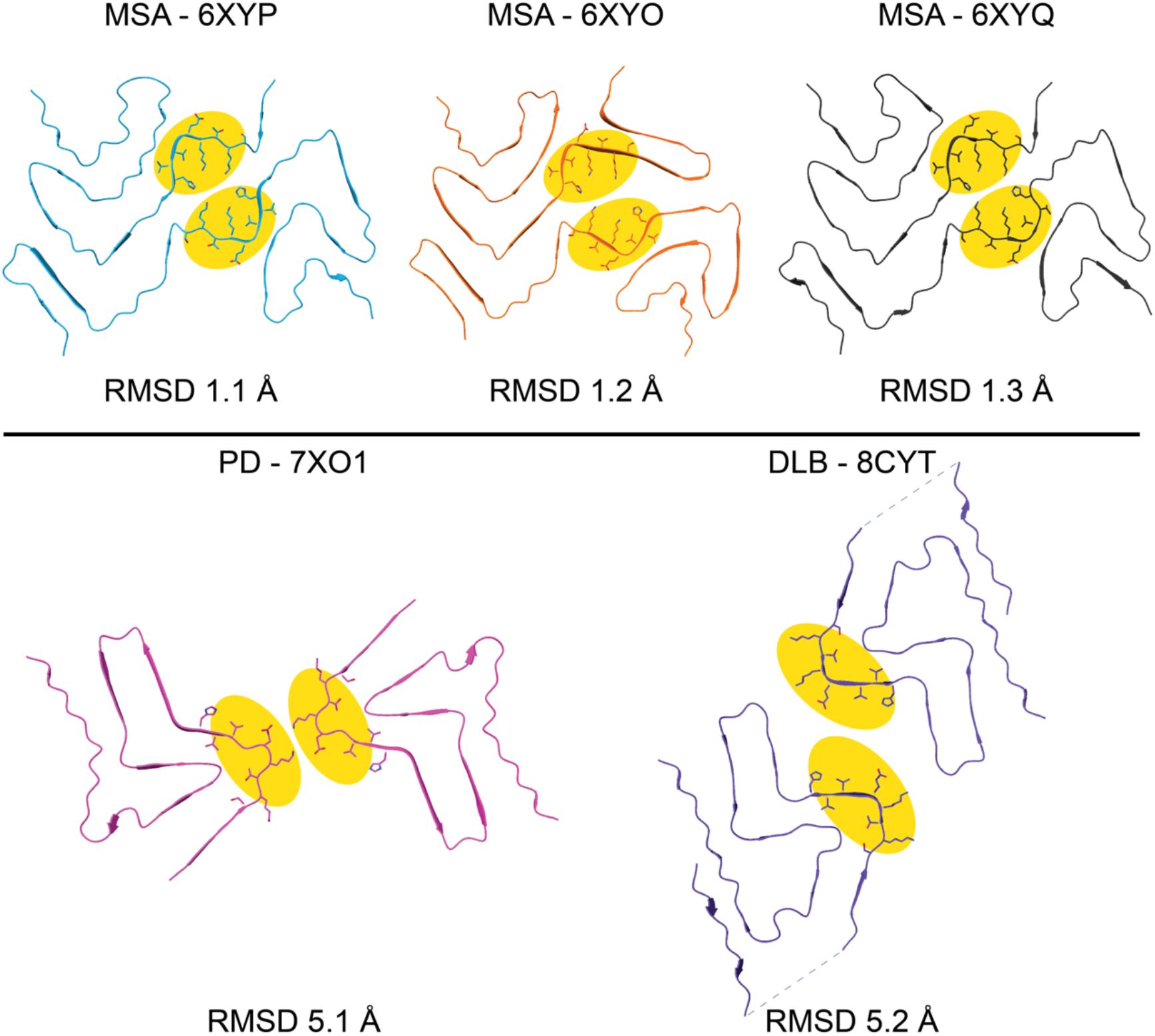
Comparison of EX-6 Bound Cryo-EM Structure to MSA, PD, and DLB Fibril Structures. Comparison of the Site 2* region (residues G41 to H50) in the EX-6 bound structure to the same area in *ex vivo* structures of αSyn fibrils provides a mechanistic explanation for the observed MSA fibril (6XYP, 6XYO, and 6XYQ) binding selectivity. In contrast, comparison of Site 2* to PD (7XO1) or DLB (8CYT) has a lower RMSD.

In order to further study [^3^H]EX-6 binding in more native environments, we performed autoradiography studies with brain tissue slices from MSA, PD, and DLB patients, as well as AD patients and healthy controls (DLB and MSA-P in **Figure 4B**, others in **Figure S18-S19**). We found high levels of binding in both MSA-C (cerebellar type) and MSA-P (Parkinson type) cases, but low binding in PD and DLB cases (**Figure 4C**), despite high levels of αSyn pathology, as determined by immunohistochemistry (IHC). Indeed, the binding that was observed in PD and DLB cases did not correlate with the severity of pathology nor its localization by IHC (**Figure 4B** and **Figure S18-S20**). In contrast, the locations of high [^3^H]EX-6 signal in MSA cases correlated well with the IHC signal for pathological αSyn. Moreover, little binding was observed in an AD case with high pathology scores for both Amyloid-? and tau, supporting the selectivity of EX-6 for αSyn fibrils over other amyloids (**Figure S20**). No binding was observed in the healthy control sample. Taken together with the saturation binding data, the tissue autoradiography data strongly support our structural model for selective MSA binding based on the similarity of the well-defined Site 2* to the same region in *ex vivo* MSA patient fibril structures.

### Comparison to other published ligand-bound αSyn structures

Recent structural studies have investigated how small molecules, such as potential PET radioligands and classical amyloid dyes like thioflavin T (ThT), bind to α-synuclein fibrils. These studies aim to provide the necessary mechanistic information for further improvement in both binding affinity and fibril specificity. Tao *et al*. provided insight into how classical dyes bind to αSyn fibrils (45). Three binding mechanisms were proposed based on the analysis of their cryo-EM data; small molecules can either bind in a vertical, horizontal, or diagonal orientation relative to the fibril axis, with the weakest binding mechanism hypothesized to be the vertical orientation and the strongest being diagonal, based on the binding affinities of the molecules (45). It was proposed that the diagonal and horizontal orientations were more favorable because inter-ligand π-stacking interactions helped to stabilize bound conformations further. Our structural analysis clearly indicates that EX-6 binds in a horizontal orientation at Site 2*. However, our novel observation of alternating ligand conformations, specifically the rotations of the fluorinated ring and the dimethyl-benzene ring, allow for stabilizing ligand-ligand π-stacking interactions, similar to the favorable π-stacking interactions observed in the Tao *et al*. diagonal orientation (45). Following on this work, Xiang *et al*. assessed PET radioligand [^18^F]-F0502B in complex with *in vitro* αSyn fibrils and observed a diagonal stacking of ligands along the fibril axis (46). Interestingly, binding of F0502B occurs at a pocket comprised of Y39 to G47 and K80-E83. This is similar to the original Site 2 identified in our computational analysis of the 2N0A ssNMR structure, where a twist of the backbone places Y39 on the opposite face of the ?-strand from H50 (**Figure S1**). We suspect that this region of the fibril is organized differently due to the presence of phosphate in our fibrilization buffer. As previously discussed, these ions enable K43, K45, and K58 to approach each other, followed by the folding back of the N-terminus between residues V15 and T22. However, in the analysis of both EX-6 and F0502B, it appears that inter-ligand π-stacking interactions and ligand-protein π-stacking at Y39 are crucial. In addition to the structural information from recent studies, multi-site ligand binding with stacked molecules along the fibril axis was reported through various computational and biochemical experiments. Using surface plasmon resonance, Sobek et al. showed that a simple kinetic model for binding of aromatic molecules with broad amyloid affinity to αSyn fibrils did not fit the data as well as a multiple binding event model (47). Furthermore, it has been demonstrated that utilizing modified molecules to promote the self-association of aromatic ThT-like ligands increases their affinity and Hill coefficient when complexed with amyloid fibrils (48). These findings have spurred the development of computational tools that account for the stacking of aromatic ligands in amyloid binding cavities (49). While our findings are consistent with those of other structural and binding studies, the alternating conformations of EX-6 in our structure provide an additional consideration for ligand optimization.

## Conclusion

The development of PET ligands for imaging αSyn fibrils, particularly those with selectivity among the PD/DLB and MSA fibril polymorphs, has the potential to provide a molecular basis for diagnosis and a means of monitoring the efficacy of investigational therapies. Our previous work, using an ultra-high-throughput *in silico* screen, identified EX-6 as a lead compound in the development of PET radiotracers for synucleinopathies (33). Here, we have performed a multimodal investigation of the binding of EX-6 to *in vitro* fibrils that recapitulate many of the structural features of MSA fibrils extracted from patients, particularly in the region referred to as Site 2* (Y39 to H50). From our cryo-EM, it is apparent that the ligand binds primarily at Site 2* at high ligand:protein concentrations. The cryo-EM structure has the most refined density at Site 2* owing to high occupancy at this site. In our XL-MS experiments, additional sites are identified; however, at low ligand concentrations, the tryptic peptide corresponding to Site 2* is the most abundant. The primacy of Site 2* is consistent with the high levels of MSA fibril selectivity observed in saturation binding studies and autoradiography.

The systematic set of structural, chemical biology, and radiography experiments conducted here has furthered that development and provided a rational basis for focusing on EX-6 derivatives as MSA imaging probes. Cryo-EM structural studies identified a set of cation-π interactions between the positively charged lysine residues (K45, K58) of αSyn and the electron-rich face of the dimethyl-benzene ring of the EX-6 ligand that stabilize the ligand near the fibril core. Previous work has shown that cation-π interactions can be as long as 6 Å and both lysine residues satisfy this requirement (50). Adjacent dimethyl-benzene rings are also stabilized by offset π-stacking interactions. Towards the solvent space, the alternating conformations of the fluorinated rings allow for a network of ligand-ligand and ligand-protein π-stacking interactions that stabilize binding at this pocket. Both offset and edge-to-face π-system interactions are observed, with favorable orientations based on the electrostatic potential surfaces of the fluorobenzene and Y39 rings.These key insights should provide a basis for rational ligand design and for optimizing the current set of Exemplar-based lead compounds already identified (33) as well as other compounds under investigation with MSA selectivity (51, 52).

### Limitations

One aspect of our study that has not been fully explored is the use of *in vitro* fibrils and not patient-derived fibrils for this structural analysis. Here, we aimed to show a multi-modal approach for determining and validating ligand binding to fibrils by conducting parallel XL-MS, FRET, and cryo-EM studies. To conduct such an investigation, a large sample volume was necessary, and this was best accomplished by *in vitro* preparations. Unfortunately, for patient-derived fibril preparations reported to date, there is simply not enough biological material extracted for this multi-modal approach. While templated amplification from MSA patient fibrils is an appealing strategy, these approaches do not appear to replicate the folds observed in patients (53, 54). In fact, the resulting polymorphs resemble *ex vivo* MSA fibrils less than *in vitro* polymorph studied here. Further studies on the specific binding to this site and identification of other possible ligand binding sites in patient-derived fibrils are the focus of our ongoing ligand-binding studies of the Exemplar-derived molecules.

Finally, the optimization of *in vivo* PET imaging ligands will need to focus on improving molecule solubility, their ability to penetrate the blood-brain barrier, and enhancing their selectivity for αSyn over A? and tau fibrils. Some of this optimization will use empirical knowledge of the pharmacological-kinetic properties of various molecular functional groups. Other aspects of this optimization will require experimental data, in the form of additional structural studies, to fully understand the physical properties that drive ligand-fibril binding. Together with computational modeling, these data can be used to systematically optimize small molecules for use as PET radioligands in the clinic.

## Materials and Methods

### Fibril formation

αSyn fibrilization experiments were conducted by incubating triply labeled (^13^C, ^2^H, ^15^N) *∝*Syn monomers in fibrilization buffer (50 mM sodium phosphate, 0.1 mM EDTA, 0.02% sodium azide, pH 7.4) for 3 weeks at 37 °C. After 3 weeks, the EX-6 ligand was added to a set of fibrils in a 1:2 molar ratio of protein to ligand and allowed to incubate for an additional 3 weeks. In all, both the apo and ligand-bound fibrils were allowed to incubate for 6 weeks before structural analysis. Fibrils for FRET were prepared similarly, and XL-MS comparisons with samples used in cryo-EM were used to validate the consistency of fibril morphologies.

### Negative stain transmission electron microscopy

The initial *∝*Syn protein concentration for TEM was 13 mg/mL. Concentrated samples, along with several dilutions, were prepared for NS-TEM. Four μL of each dilution were applied to glow-discharged 200-mesh copper grids with a continuous carbon film and stained with 1% uranyl acetate for 10 seconds before blotting with filter paper (Whatman grade 1). Grids were allowed to dry overnight, followed by imaging on a Talos L120C operated at 120 kV. Images were collected at a pixel size of 2.59 Å with a total dose of 25 e^-^/Å^2^. Images were analyzed using the IMOD software suite (55).

### Cryo-EM grid preparation and data collection

Cryo-EM grids were prepared by applying 4 μL of *∝*-synuclein fibrils to glow-discharged Quantifoil R2/1 200 mesh, copper grids with a wait time of 60 seconds. The grids were then plunged into a bath of liquid ethane using a Vitrobot Mark IV operating at 20°C and a relative humidity of 95%, to generate vitrified samples. Blot forces ranging from +4 to +6 with a blot time of 4 seconds and a drain time of 0.5 seconds were tested.

Data collection was performed on a Titan Krios G3i FEG-TEM operated at 300 kV, equipped with a Gatan K3 direct electron detector and BioQuantum energy filter. Dose-fractionated micrographs were collected in correlated double sampling (CDS), counting mode at a pixel size of 0.834 Å over a defocus range of -0.5 to -2.5 μm in increments of 0.25 μm. Movies were acquired with a total dose of 40 e^-^/Å^2^ with three shots per hole, acquiring multiple holes per stage movement by using EPU/AFIS. On average, ∼300 movies were collected per hour.

### Cryo-EM helical reconstruction

Helical reconstruction data processing was performed within the RELION-4.0 framework (56) and extensively detailed in Sanchez et al. (36). Briefly, preprocessing was performed on gain-normalized micrographs, which comprised 40 frames. The frames were motion corrected with MotionCor2, and initial estimates for whole-micrograph contrast transfer function (CTF) values were performed with gCTF (57, 58). Fibrils were picked using the Topaz-filament executable (59, 60). Selected segments were subjected to particle extraction with a box size of 864 pixels, followed by 2D classification to determine the cross-over distance. An initial model was generated using the *relion_helix_inimodel2d* program (61, 62). Several rounds of 2D classification, 3D classification, and 3D auto-refinement were required to determine the optimal helical parameters and final particle set. After the final round of 3D auto-refinement, a soft-edge solvent mask comprising 25% of the central Z height was applied to the map, followed by sharpening using standard post-processing in RELION. The *relion_helix_toolbox* program was used to apply real-space symmetry to the final map (61). Finally, a local resolution map was generated using the automated post-processing job. A similar processing scheme was followed for the EX-6 ligand-fibril complex.

### Model building, ligand generation and docking, and validation

A molecular model of the apo fibril structure was built into the final map by first docking in a previously solved cryo-EM structure (PDB: 6H6B) (22). Extensive rebuilding of the N-terminal region was performed in COOT due to the presence of an additional density not observed in previous *in vitro* a-syn structures (63, 64). Residues 15-22 and 36-97 were resolvable in our cryo-EM map. The structure was then refined in PHENIX with rotamer, Ramachandran restraints, and non-crystallographic symmetry (NCS) imposed (65, 66). These steps were repeated for the ligand-bound structure using the refined apo structure as the initial model. Residues 15-22 and 36-96 were resolvable in the ligand-bound cryo-EM map. The protein aspect of the model was refined in Phenix similarly to the apo structure. For the EX-6 ligand, the simplified molecular-input line-entry system (SMILES) string was provided as input to the phenix.elbow program to generate initial coordinates, followed by optimization of ligand geometry using the eLBOW AM1 QM method (67). The ligand was then manually docked into the cryo-EM density using ChimeraX. Lastly, refinement of the protein and ligand structure was performed in PHENIX with rotamer and Ramachandran restraints for the protein, and ligand geometry restraints using the eLBOW restraints file. The final models displayed excellent validation statistics, as reported in PHENIX (66).

### Structure analysis

Structural analysis was performed using ChimeraX for visualization of electrostatic potentials and hydrophobicity of the model (68, 69). Buried surface area calculations and molecular interactions were analyzed using the Proteins, Interfaces, Structures, and Assemblies (PISA) server (38, 39).

### XL-MS Sample Preparation and Analysis

Samples from cryo-EM were pelleted at 16.1 RPM for 90 minutes, washed 3 times with PBS containing 0.02% sodium azide, and reconstituted in PBS with 0.02% sodium azide. The sample mixture was then vigorously shaken and vortexed to resuspend the pellet. A sample of the resuspended fibril solution was then subjected to boiling in 3.6% SDS, followed by a DC assay. After quantification samples were made containing either 1 µM fibrils at 1000 µL solution volume with 500 nM EX-6-CLX in PBS or 10 µM fibrils at 100 µL with 500 nM EX-6-CLX .DMSO content was kept constant at 1% for all solutions. After 1 hour of incubation at 37 °C, the samples were irradiated at 365 nm in 1 mL glass tubes using a Lumidox array for 2 minutes at 355 mW/cm^2^. The samples were then transferred to 1.5 mL Eppendorf tubes, frozen in liquid nitrogen, and the solution was dried under vacuum. After drying, samples were reconstituted in 100 µL water and subjected to chloroform/methanol precipitation. To the dried solutions, 20 µL of ammonium bicarbonate buffer was added along with 0.36 µg trypsin (1:20 enzyme:protien). After 2 hours incubation, the sample was acidified to 0.1% trifluoroacetic acid (TFA) total concentration using 10% TFA. Alkylated dihydroxybenzoic acid synthesized as previously described and disolved in 50:50 acetonitrile/water with 0.1% TFA to a concentration of 5 mg/mL (70). This solution was added 1:10 to a solution of 10 mg/mL α-cyano-4-hydroxycinnamic acid in 50:50 acetonitrile/water with 0.1% TFA to create the final matrix solution. One part matrix solution and one part sample solution were mixed and deposited on a ground steel MALDI plate from bruker.

Spectra were acquired on a Rapiflex MALDI-TOF from Bruker in reflector positive mode from 700-3200 Da at 60% laser power. Calibration was performed using Peptide Calibration Standard II from Bruker and applied to adjacent sample spots. Each spectrum had its baseline subtracted and smoothed in FlexAnalysis. Following normalization, a mass table was generated. This table was then transferred to BioTools where the mass table was compared to that of a theoretical digest containing potential crosslinked peptides. Peaks that corresponded to the crosslinked mass were then identified and subject to MS2 analysis to confirm crosslinking. Further analysis of these spectra is supplied in the Supporting Information.

## Acknowledgments

This work was supported in part by the University of Wisconsin-Madison, and public health service grants U24 GM139168 to E.R.W., P41 GM136463 to C.M.R., RF1 NS103873 to E.J.P., U19 NS110456 to E.J.P. and R.M.H., and RF1 NS110436 to E.R.W., C.M.R., and P.T.K. from the NIH.J.C.S. was supported in part by the Biotechnology Training Program at the University of Wisconsin, Madison, T32 GM135066, the Steenbock Predoctoral Graduate Fellowship administered by the University of Wisconsin-Madison Department of Biochemistry, and the SciMed Graduate Research Scholars Fellowship with support for this fellowship provided by the Graduate School, part of the Office of Vice Chancellor for Research and Graduate Education at the University of Wisconsin-Madison, with funding from the Wisconsin Alumni Research Foundation and the UW-Madison.C.G.B. was supported by the NIH Ruth L. Kirschstein Fellowship, F32 GM149118, from the NIGMS.R.M.P. was supported by the NIH Chemistry Biology Interface Training Program (NIH T32-GM133398) and by an individual F31 fellowship. Instruments supported by the NIH and NSF include NMR (NSF CHE 1827457) and mass spectrometers (NIH S10 OD030460)

## References

1. S. Koga, H. Sekiya, N. Kondru, O. A. Ross, D. W. Dickson, Neuropathology and molecular diagnosis of Synucleinopathies. Molecular Neurodegeneration 16, 83 (2021).

2. C. H. Adler et al., Clinical Diagnostic Accuracy of Early/Advanced Parkinson Disease. Neurology Clinical Practice 11, e414–e421 (2021).

3. J. Jankovic, A. H. Rajput, M. P. McDermott, D. P. Perl, G. for the Parkinson Study, The Evolution of Diagnosis in Early Parkinson Disease. Archives of Neurology 57, 369–372 (2000).

4. R. Qi et al., A blood-based marker of mitochondrial DNA damage in Parkinson’s disease. Science Translational Medicine 15, eabo1557 (2023).

5. A. Siderowf et al., Assessment of heterogeneity among participants in the Parkinson’s Progression Markers Initiative cohort using alpha-synuclein seed amplification: a cross-sectional study. The Lancet Neurology 22, 407–417 (2023).

6. C. H. Gibbons et al., Skin Biopsy Detection of Phosphorylated α-Synuclein in Patients With Synucleinopathies. JAMA 331, 1298–1306 (2024).

7. H. I. Hurtig et al., Alpha-synuclein cortical Lewy bodies correlate with dementia in Parkinson’s disease. Neurology 54, 1916–1921 (2000).

8. R. L. Miller et al., Quantifying regional alpha-synuclein, amyloid beta, and tau accumulation in lewy body dementia. Ann Clin Transl Neurol 9, 106–121 (2022).

9. M. A. Hely, W. G. J. Reid, M. A. Adena, G. M. Halliday, J. G. L. Morris, The Sydney multicenter study of Parkinson’s disease: The inevitability of dementia at 20 years. Movement Disorders 23, 837–844 (2008).

10. R. Jakes, M. G. Spillantini, M. Goedert, Identification of two distinct synucleins from human brain. FEBS letters 345, 27–32 (1994).

11. W. S. Davidson, A. Jonas, D. F. Clayton, J. M. George, Stabilization of alpha-synuclein secondary structure upon binding to synthetic membranes. J Biol Chem 273, 9443–9449 (1998).

12. A. Iwai et al., The precursor protein of non-Aβ component of Alzheimer’s disease amyloid is a presynaptic protein of the central nervous system. Neuron 14, 467–475 (1995).

13. H. T. Li, H. N. Du, L. Tang, J. Hu, H. Y. Hu, Structural transformation and aggregation of human α‐synuclein in trifluoroethanol: Non‐amyloid component sequence is essential and β‐sheet formation is prerequisite to aggregation. Biopolymers: Original Research on Biomolecules 64, 221–226 (2002).

14. H.-J. Lee, C. Choi, S.-J. Lee, Membrane-bound α-synuclein has a high aggregation propensity and the ability to seed the aggregation of the cytosolic form. Journal of Biological Chemistry 277, 671–678 (2002).

15. R. A. Crowther, R. Jakes, M. G. Spillantini, M. Goedert, Synthetic filaments assembled from C-terminally truncated α-synuclein. FEBS letters 436, 309–312 (1998).

16. S. X. Pancoe et al., Effects of Mutations and Post-Translational Modifications on α-Synuclein In Vitro Aggregation. Journal of Molecular Biology 434, 167859 (2022).

17. H. A. Lashuel, C. R. Overk, A. Oueslati, E. Masliah, The many faces of α-synuclein: from structure and toxicity to therapeutic target. Nature Reviews Neuroscience 14, 38–48 (2013).

18. R. Khounlo et al., Membrane Binding of α-Synuclein Stimulates Expansion of SNARE-Dependent Fusion Pore. Frontiers in Cell and Developmental Biology 9 (2021).

19. T. Bartels, J. G. Choi, D. J. Selkoe, α-Synuclein occurs physiologically as a helically folded tetramer that resists aggregation. Nature 477, 107–110 (2011).

20. W. Wang et al., A soluble α-synuclein construct forms a dynamic tetramer. Proceedings of the National Academy of Sciences 108, 17797–17802 (2011).

21. M. G. Spillantini, R. A. Crowther, R. Jakes, M. Hasegawa, M. Goedert, alpha-Synuclein in filamentous inclusions of Lewy bodies from Parkinson’s disease and dementia with lewy bodies. Proc Natl Acad Sci U S A 95, 6469–6473 (1998).

22. R. Guerrero-Ferreira et al., Cryo-EM structure of alpha-synuclein fibrils. Elife 7 (2018).

23. Y. Li et al., Amyloid fibril structure of alpha-synuclein determined by cryo-electron microscopy. Cell Res 28, 897–903 (2018).

24. D. R. Boyer et al., Structures of fibrils formed by α-synuclein hereditary disease mutant H50Q reveal new polymorphs. Nature structural & molecular biology 26, 1044–1052 (2019).

25. K. Zhao et al., Parkinson’s disease associated mutation E46K of α-synuclein triggers the formation of a distinct fibril structure. Nature communications 11, 2643 (2020).

26. Y. Sun et al., Cryo-EM structure of full-length α-synuclein amyloid fibril with Parkinson’s disease familial A53T mutation. Cell research 30, 360–362 (2020).

27. M. Baba et al., Aggregation of alpha-synuclein in Lewy bodies of sporadic Parkinson’s disease and dementia with Lewy bodies. Am J Pathol 152, 879–884 (1998).

28. Y. Yang et al., New SNCA mutation and structures of alpha-synuclein filaments from juvenile-onset synucleinopathy. Acta Neuropathol 145, 561–572 (2023).

29. Y. Yang et al., Structures of α-synuclein filaments from human brains with Lewy pathology. Nature 610, 791–795 (2022).

30. M. Schweighauser et al., Structures of α-synuclein filaments from multiple system atrophy. Nature 585, 464–469 (2020).

31. M. M. Alam, S. H. Lee, S. Wasim, S.-Y. Lee, PET tracer development for imaging α-synucleinopathies. Archives of Pharmacal Research 48, 333–350 (2025).

32. J. Therriault et al., Biomarker-based staging of Alzheimer disease: rationale and clinical applications. Nature Reviews Neurology 20, 232–244 (2024).

33. J. J. Ferrie et al., Identification of a nanomolar affinity α-synuclein fibril imaging probe by ultra-high throughput in silico screening. Chem Sci 11, 12746–12754 (2020).

34. M. D. Tuttle et al., Solid-state NMR structure of a pathogenic fibril of full-length human alpha-synuclein. Nat Struct Mol Biol 23, 409–415 (2016).

35. M. H. Milchberg et al., Alpha-Synuclein Fibril Structures Cluster into Distinct Classes. bioRxiv 10.1101/2025.04.30.651534, 2025.2004.2030.651534 (2025).

36. J. C. Sanchez, J. A. Pierson, C. G. Borcik, C. M. Rienstra, E. R. Wright, High-resolution Cryo-EM Structure Determination of a-Synuclein—A Prototypical Amyloid Fibril. Bioprotocol 15, e5171 (2025).

37. M. B. Cory et al., FRETing about the details: Case studies in the use of a genetically encoded fluorescent amino acid for distance-dependent energy transfer. Protein Science 32, e4633 (2023).

38. E. Krissinel, Stock-based detection of protein oligomeric states in jsPISA. Nucleic Acids Research 43, W314–W319 (2015).

39. E. Krissinel, Protein interfaces, surfaces and assemblies service PISA at European Bioinformatics Institute. J Mol Biol 372, 774 (2007).

40. M. R. Sawaya, M. P. Hughes, J. A. Rodriguez, R. Riek, D. S. Eisenberg, The expanding amyloid family: Structure, stability, function, and pathogenesis. Cell 184, 4857–4873 (2021).

41. I. Sungwienwong et al., Improving target amino acid selectivity in a permissive aminoacyl tRNA synthetase through counter-selection. Organic & Biomolecular Chemistry 15, 3603–3610 (2017).

42. C. M. Haney et al., Comparison of strategies for non-perturbing labeling of α-synuclein to study amyloidogenesis. Organic & Biomolecular Chemistry 14, 1584–1592 (2016).

43. A. V. West et al., Labeling Preferences of Diazirines with Protein Biomolecules. Journal of the American Chemical Society 143, 6691–6700 (2021).

44. Y. Jiang et al., Dissecting diazirine photo-reaction mechanism for protein residue-specific cross-linking and distance mapping. Nature Communications 15, 6060 (2024).

45. Y. Tao et al., Structural mechanism for specific binding of chemical compounds to amyloid fibrils. Nature Chemical Biology 19, 1235–1245 (2023).

46. J. Xiang et al., Development of an alpha-synuclein positron emission tomography tracer for imaging synucleinopathies. Cell 186, 3350-3367.e3319 (2023).

47. J. Sobek et al., Efficient characterization of multiple binding sites of small molecule imaging ligands on amyloid-beta, tau and alpha-synuclein. European Journal of Nuclear Medicine and Molecular Imaging 51, 3960–3977 (2024).

48. J. L. Cifelli, C. C. Capule, J. Yang, Noncovalent, Electrostatic Interactions Induce Positively Cooperative Binding of Small Molecules to Alzheimer’s and Parkinson’s Disease-Related Amyloids. ACS Chemical Neuroscience 10, 991–995 (2019).

49. M. S. Smith et al., Docking for Molecules That Bind in a Symmetric Stack with SymDOCK. Journal of Chemical Information and Modeling 64, 425–434 (2024).

50. J. P. Gallivan, D. A. Dougherty, Cation-π interactions in structural biology. Proceedings of the National Academy of Sciences 96, 9459–9464 (1999).

51. H. Y. Kim et al., A Novel Brain PET Radiotracer for Imaging Alpha Synuclein Fibrils in Multiple System Atrophy. Journal of Medicinal Chemistry 66, 12185–12202 (2023).

52. Z. Lengyel-Zhand et al., Synthesis and characterization of high affinity fluorogenic α-synuclein probes. Chemical Communications 56, 3567–3570 (2020).

53. S. Lövestam et al., Seeded assembly in vitro does not replicate the structures of α-synuclein filaments from multiple system atrophy. FEBS Open Bio 11, 999–1013 (2021).

54. Y. Fichou et al., Cofactors are essential constituents of stable and seeding-active tau fibrils.Proceedings of the National Academy of Sciences 115, 13234–13239 (2018).

55. J. R. Kremer, D. N. Mastronarde, J. R. McIntosh, Computer Visualization of Three-Dimensional Image Data Using IMOD. Journal of Structural Biology 116, 71–76 (1996).

56. D. Kimanius, L. Dong, G. Sharov, T. Nakane, S. H. W. Scheres, New tools for automated cryo-EM single-particle analysis in RELION-4.0. Biochemical Journal 478, 4169–4185 (2021).

57. K. Zhang, Gctf: Real-time CTF determination and correction. Journal of Structural Biology 193, 1–12 (2016).

58. S. Q. Zheng et al., MotionCor2: anisotropic correction of beam-induced motion for improved cryo-electron microscopy. Nat Methods 14, 331–332 (2017).

59. T. Bepler et al., Positive-unlabeled convolutional neural networks for particle picking in cryo-electron micrographs. Nat Methods 16, 1153–1160 (2019).

60. S. Lovestam, S. H. W. Scheres, High-throughput cryo-EM structure determination of amyloids. Faraday Discuss 240, 243–260 (2022).

61. S. He, S. H. W. Scheres, Helical reconstruction in RELION. J Struct Biol 198, 163–176 (2017).

62. S. H. W. Scheres, Amyloid structure determination in RELION-3.1. Acta Crystallogr D Struct Biol 76, 94–101 (2020).

63. P. Emsley, B. Lohkamp, W. G. Scott, K. Cowtan, Features and development of Coot. Acta Crystallogr D 66, 486–501 (2010).

64. P. Emsley, K. Cowtan, Coot: model-building tools for molecular graphics. Acta crystallographica section D: biological crystallography 60, 2126–2132 (2004).

65. D. Liebschner et al., Macromolecular structure determination using X-rays, neutrons and electrons: recent developments in Phenix. Acta Crystallogr D Struct Biol 75, 861–877 (2019).

66. P. V. Afonine et al., Real-space refinement in PHENIX for cryo-EM and crystallography. Acta Crystallographica Section D: Structural Biology 74, 531–544 (2018).

67. N. W. Moriarty, R. W. Grosse-Kunstleve, P. D. Adams, electronic Ligand Builder and Optimization Workbench (eLBOW): a tool for ligand coordinate and restraint generation. Acta Crystallographica Section D 65, 1074–1080 (2009).

68. E. C. Meng et al., UCSF ChimeraX: Tools for structure building and analysis. Protein Sci 32, e4792 (2023).

69. E. F. Pettersen et al., UCSF ChimeraX: Structure visualization for researchers, educators, and developers. Protein Sci 30, 70–82 (2021).

70. Y. Fukuyama et al., Alkylated Dihydroxybenzoic Acid as a MALDI Matrix Additive for Hydrophobic Peptide Analysis. Analytical Chemistry 84, 4237–4243 (2012).

