## Supplementary Materials for "Complementary Structural and Chemical Biology Methods Reveal the Basis for Selective Radioligand Binding to α-Synuclein in MSA Tissue"

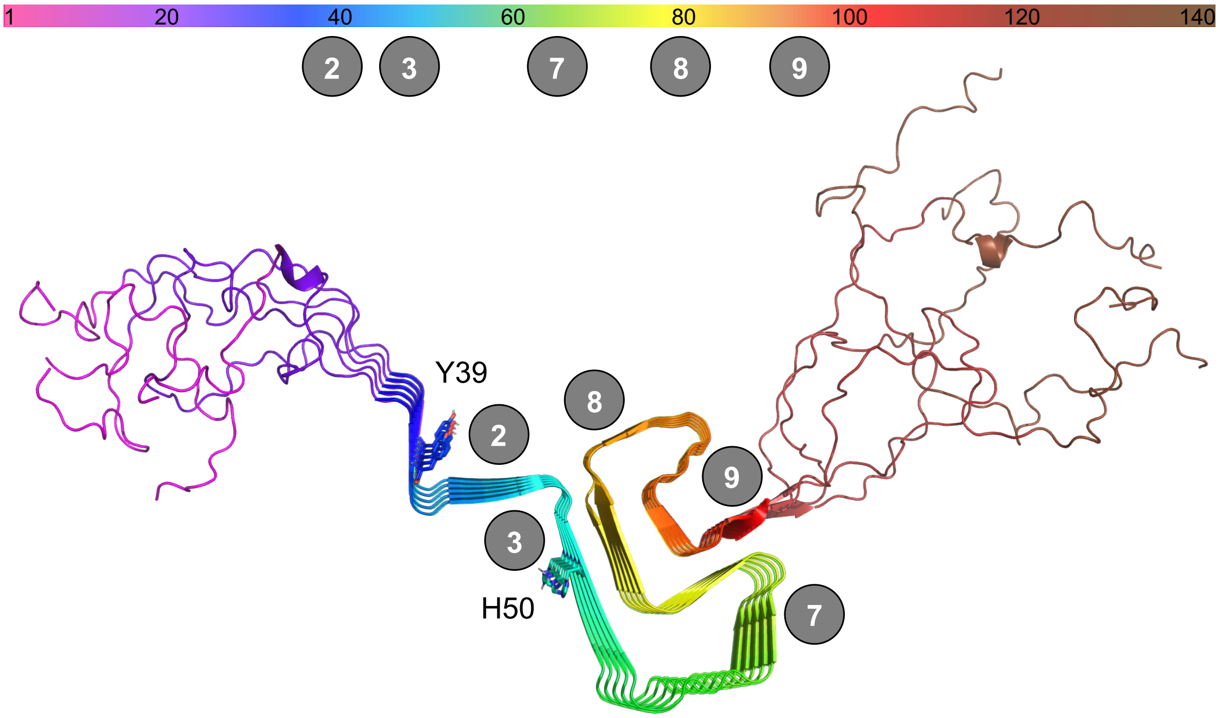


**Figure S1** Potential small molecule binding sites on αSyn fibrils identified using PDB ID 2N0A (1) in Hsieh et al. (2).

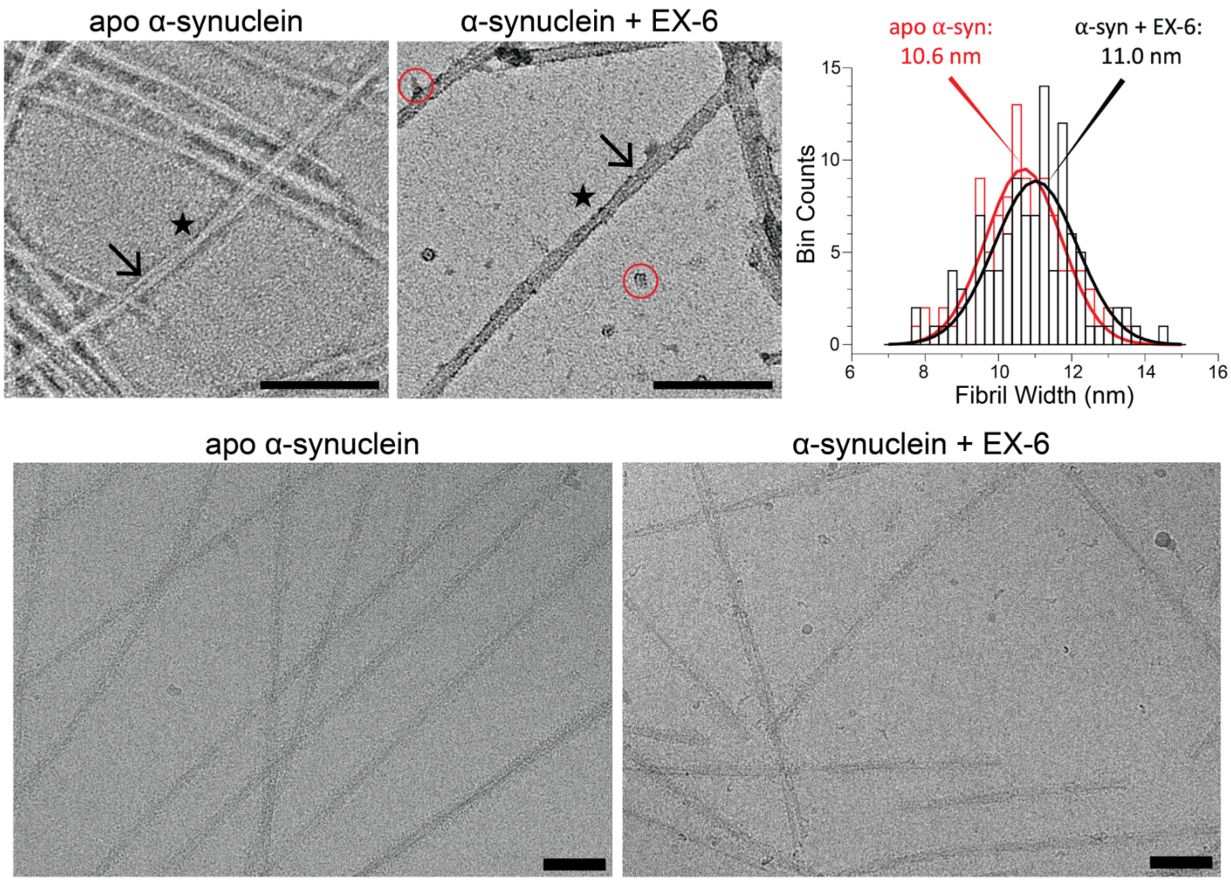


**Figure S2** Comparison of αSyn Fibrils with and Without EX-6 Reveals No Major Changes to Fibrils. Top Left: NS-TEM images of αSyn fibrils stained with 1% uranyl acetate. The arrow denotes the region where the two protofilaments are discernible. An asterisk denotes a crossover point, indicating that fibrils are twisting. Red circles show clumping likely due to excess ligands. Scale bar, 100 nm. Top Right: Fibril width measurements from NS-TEM images of apo and EX-6 bound fibrils. Bottom: Cryo-EM micrographs of twisting αSyn fibrils with and without EX-6 ligands. Scale bar, 50 nm.

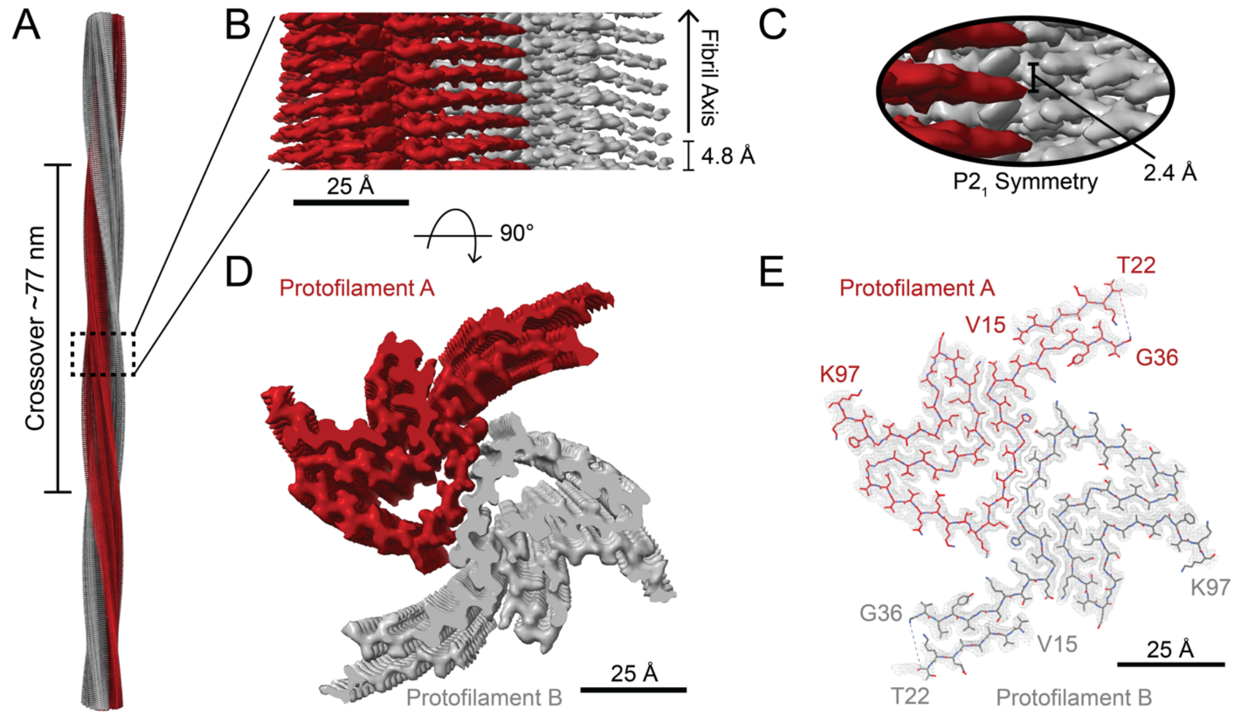


**Figure S3** Cryo-EM Data (EMD-45639) and Model (PDB: 9CK3) for αSyn (Apo) Fibrils. A. Multiple maps aligned for easy visualization of the twisting nature of the fibrils. B. Close-up of Site 2 with no ligand density. The map was segmented and colored for easy identification of each protofilament. C. The protofilaments exhibit pseudo-screw symmetry, which was applied during reconstruction. D. Cross-section of the central region of the map. E. Map-model overlay with residues V15 to T22, and G36 to K97 resolved in the map.

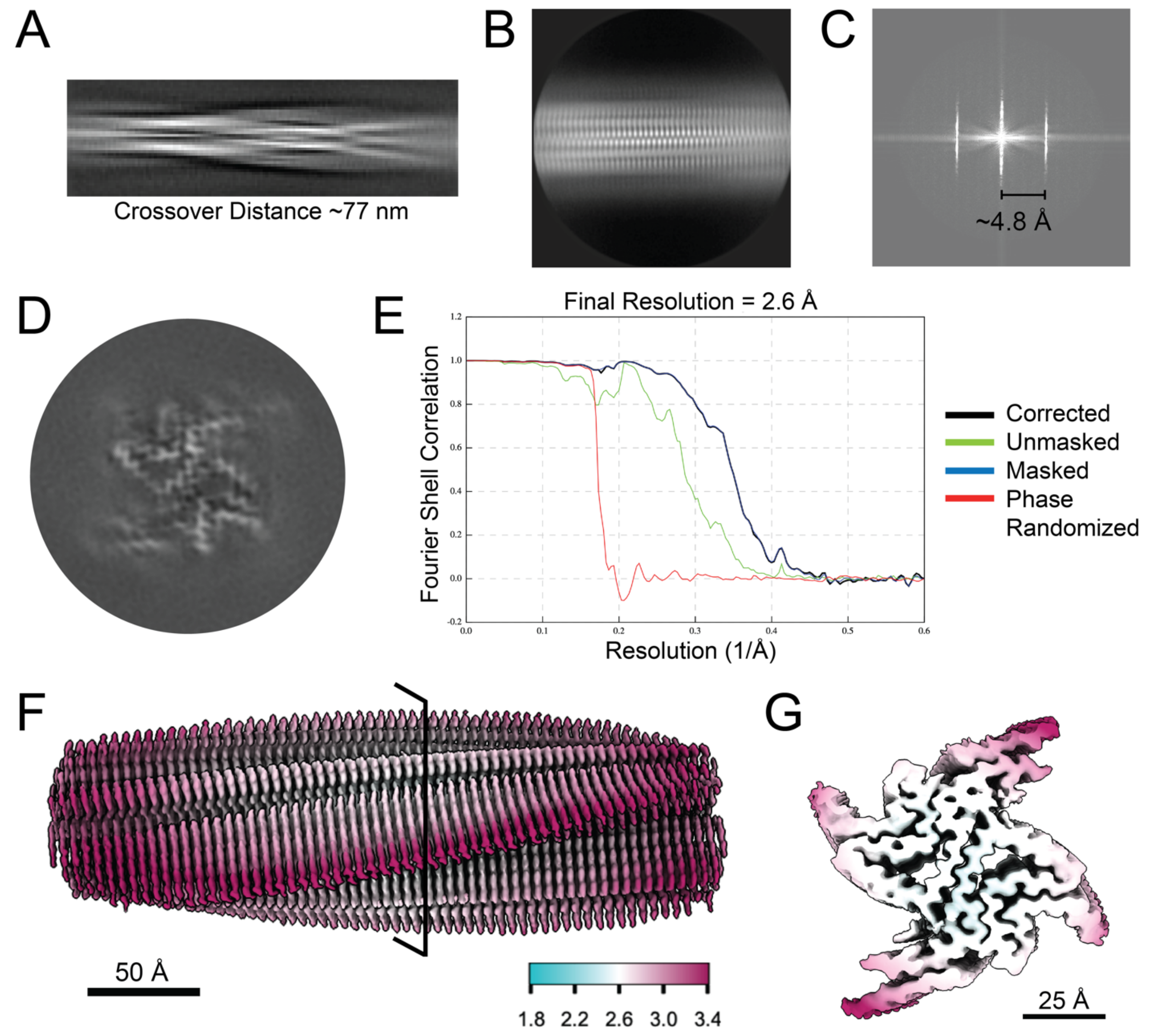


**Figure S4** Cryo-EM Data Processing Results and Local Resolution Map for Ex-6 Bound Fibrils. A. 2D class average (864-pixel box) used to estimate crossover distance. B. 2D class average (360-pixel box) used to verify helical rise. C. Average power spectrum showing a layer line at ~4.8 Å. D. Cross-section of the 3D auto-refinement showing two protofilaments with amyloid “Greek key” conformations. E. Gold-standard Fourier Shell Correlation at a resolution of 2.6 Å. F. Local resolution map of the Ex-6 bound fibril with the highest resolution towards the center of the fibril axis. G. Cross-section of the local resolution map shows the highest resolution towards the fibril core and ligand density visible at Site 2*. Bar in F represents the region along the fibril axis for the cross-section in G.

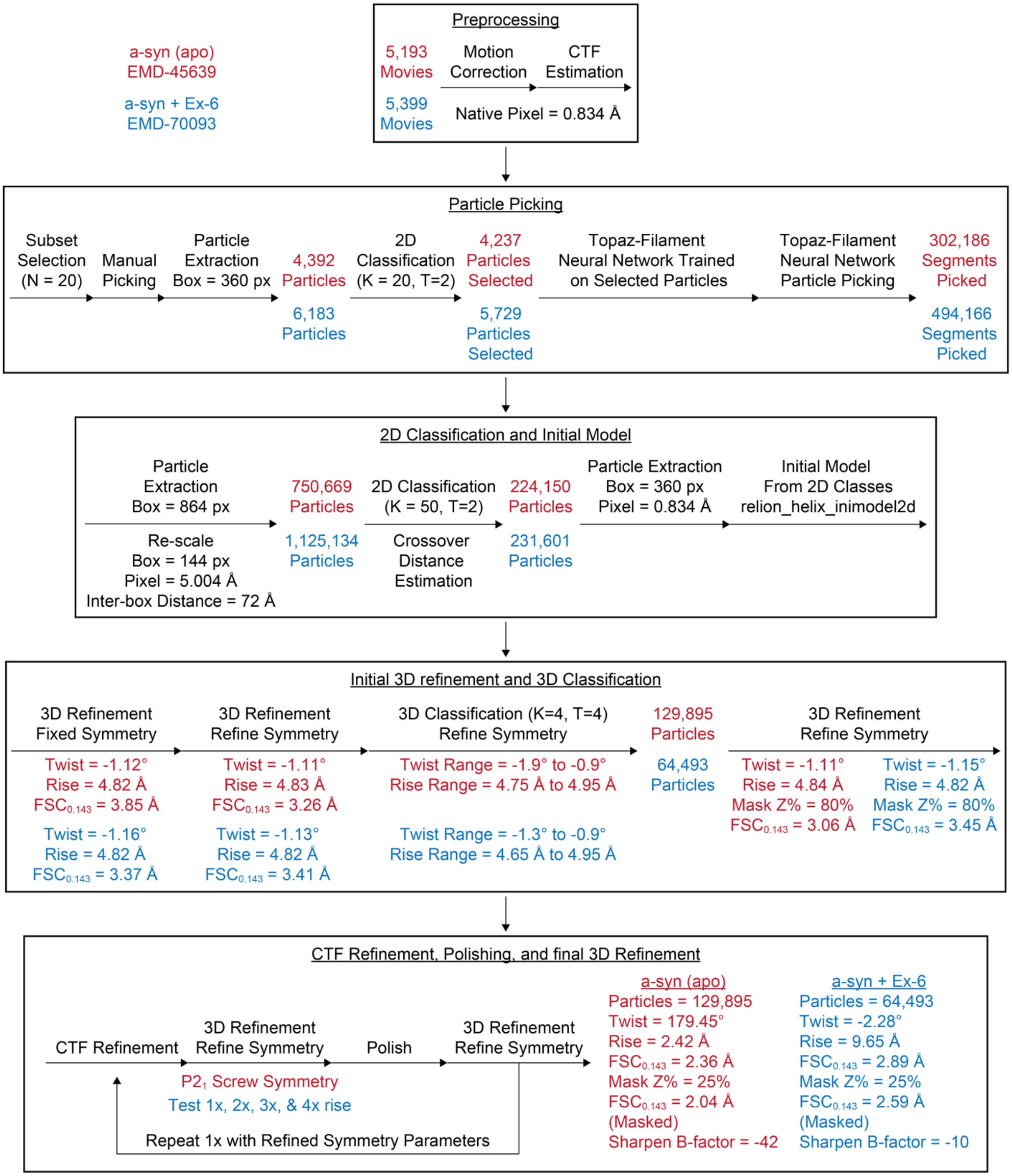


**Figure S5** Cryo-EM Helical Reconstruction Workflow. Results for each step of αSyn (red) and αSyn bound with Ex-6 (blue) processing.

**Table S1**. Statistics for Cryo-EM Data Collection, Reconstruction in RELION, and Molecular Model.

| Protein | α-syn (apo) fibril | α-syn fibril + EX-6 |
|---|---|---|
| PDB ID | 9CK3 | 9O4B |
| EMDB ID | EMD-45639 | EMD-70093 |
| **Data collection and processing** | | |
| Voltage (kV) | 300 | 300 |
| Electron exposure ($e^-/Å^2$) | 40 | 40 |
| Number of frames | 40 | 40 |
| Nominal defocus range (-μm) | 0.5 - 2.5, 0.25 | 0.5 - 2.5, 0.25 |
| Pixel size (Å) | 0.834 | 0.834 |
| Symmetry (beyond helical) | $C_1$ | $C_1$ |
| Micrographs | 5,193 | 5,399 |
| Segments | 302,186 | 494,166 |
| Initial particles | 750,669 | 1,125,134 |
| Final particles | 129,895 | 64,493 |
| Box size (Å) | 300 | 300 |
| Box size (pixels) | 360 | 360 |
| Helical symmetry | | |
| rise (Å) | 2.42 | 9.65 |
| twist (°) | 179.45 | -2.28 |
| Helical Z parameter (%) | | |
| masking | 25 | 25 |
| refinement | 25 | 25 |
| Map resolution (Å) threshold | 0.143 | 0.143 |
| Masked | 2.04 | 2.59 |
| Unmasked | 2.36 | 2.89 |
| Sharpening B factor ($Å^2$) | -43 | -10 |
| **Structure Refinement** | | |
| Total atoms | 5,748 | 5,940 |
| ligands | 0 | 12 |
| water | 0 | 0 |
| RMS, bonds (Å) | 0.004 | 0.001 |
| RMS, angles (°) | 0.643 | 0.375 |
| Ramachandran favored (%) | 98.48 | 100 |
| Ramachandran outliers (%) | 0.00 | 0.00 |
| Rotamer outliers (%) | 2.08 | 0.00 |
| Map-model correlation | 0.90 | 0.76 |
| Model B factors | | |
| minimum | 5.38 | 60.67 |
| mean | 41.28 | 101.73 |
| maximum | 88.56 | 156.72 |
| MolProbity score | 1.42 | 1.38 |
| Clashscore | 3.88 | 6.90 |

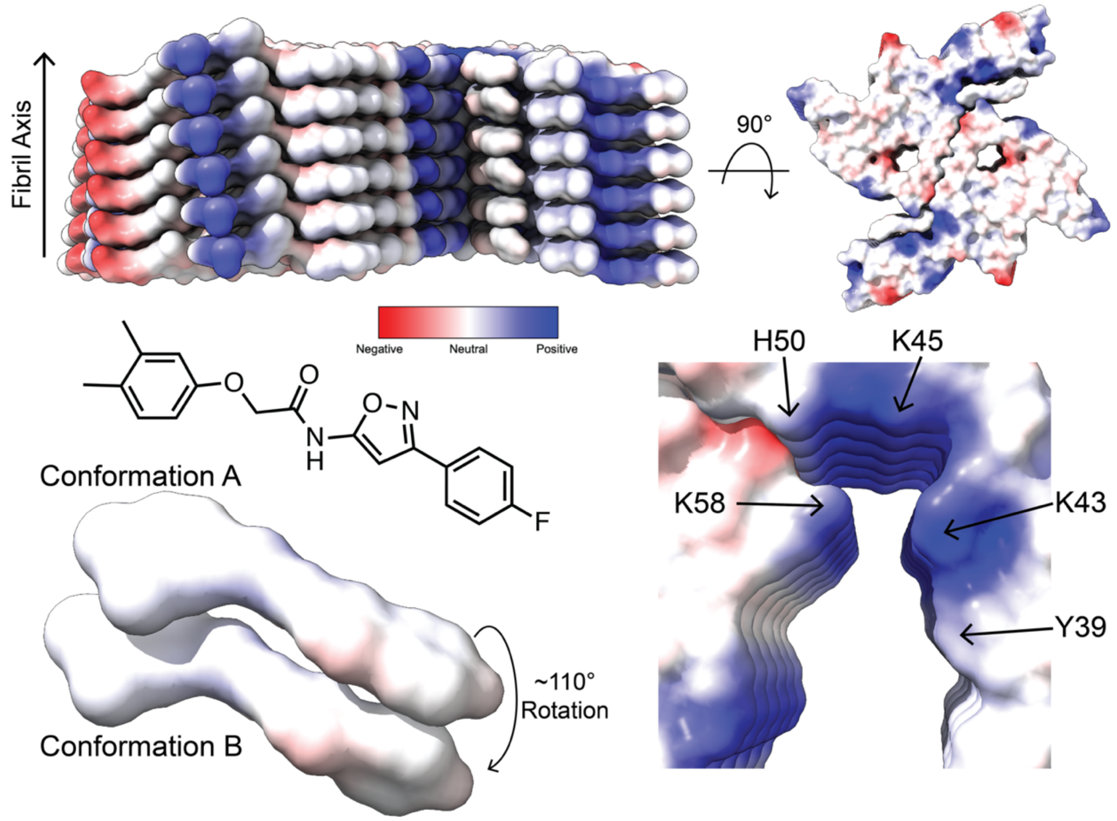


**Figure S6** Ex-6 Primarily Binds to a Surface Accessible Cationic Groove. Top: Electrostatic potential model of protein-ligand complex. Bottom Left: Ex-6 electrostatic potential model detailing the ~110° rotation of the fluorinated ring. Bottom Right: Cationic groove, denoted site 2*, comprised of Y39, K43, K45, H50, and K58.

**Helical Reconstruction Workflow**

Helical reconstruction data processing of the apo and EX-6-bound $\alpha$-synuclein fibrils was performed within the RELION-4.0 framework (**Figure S5**) (44). Preprocessing was performed on gain-normalized micrographs, which comprised 40 frames. The frames were motion-corrected using MotionCor2, and initial estimates for the whole-micrograph contrast transfer function (CTF) values were obtained with gCTF.

Fibrils were picked using the Topaz-filament executable (47, 48). Briefly, 20 micrographs were randomly selected for manual picking. This small subset was subjected to particle extraction with a box size of 360 pixels and an inter-box distance of ~40 Å, with an initial helical rise estimate of 4.82 Å. 2D classification was performed on the extracted particles using the expectation-maximization algorithm for helical segments with a T-regularization value of 2, 20 classes, and a mask diameter of 285 Å. The particles belonging to the best classes were selected as templates for training the Topaz neural network using the Topaz-filament wrapper within the RELION auto-picking job. The additional Topaz arguments -f (filament) and -t (threshold) were used for picking start-end coordinates. Additionally, thresholds between -6 to 0 were tested to determine the optimal picking parameters. Topaz picking resulted in 302,186 segments from 4,858 micrographs.

Selected segments were subjected to particle extraction with a box size of 864 pixels, rescaled to 144 pixels with a pixel size of ~5.0 Å, and an inter-box distance of 72 Å for an initial particle set of 750,669 particles. Extracted particles were used for reference-free 2D classification with a T-regularization value of 2, in-plane angular sampling from 2°, and an angular search range of +/- 18°. Next, 2D class averages were inspected and used to calculate the apparent cross-over distance of the twisting protofilaments at ~720 Å. The best 2D classes, containing 224,150 particles, were selected and used as templates for initial model generation using the *relion_helix_inimodel2d* program (49, 50). Particles pertaining to the best 2D classes were re-extracted to their original pixel size of 0.834 Å and a box size of 360 pixels. Additionally, the initial model was rescaled to the new box size and pixel size using the *relion_image_handler* program (49).

3D reconstruction required several rounds of 3D refinement. First, helical layer lines in the averaged power spectrum from the best 2D classes were analyzed to determine an initial rise estimate of 4.8 Å. The initial rise and estimated crossover distance were used to calculate an initial helical twist estimate of -1.2°. The initial model, low-pass filtered to 10 Å, and the re-extracted particles were subjected to 1 round of 3D auto-refinement with fixed initial estimates for helical twist and rise. The job resulted in a reconstruction comprised of two protofilaments with clear separation of $\beta$-strands along the helical axis, and a cross-section of the map revealed an amyloid key structural motif per protofilament.

The map and particles were then used as inputs for a round of 3D classification (k=4) with optimization of helical twist and rise values. Two classes, comprising 129,895 particles, were selected for an additional round of 3D auto-refinement with optimization of helical parameters. The resulting map displayed $P2_1$ pseudo-screw symmetry, which was applied in subsequent refinement steps. The particles were subjected to two rounds of CTF refinement and two rounds of Bayesian polishing. A 3D auto-refinement was performed after each CTF refinement and Bayesian polishing step. During these refinements, the resulting 3D reconstruction was low-pass filtered to 4.5 Å and used as the initial reference for the subsequent 3D auto-refinement job with optimization of helical parameters. Additionally, a mask comprising 80% of the central Z height was applied during these steps.

After the final round of 3D auto-refinement, soft-edge solvent masks comprising 80% and 25% of the central Z height were applied to the map, followed by sharpening using standard post-processing in RELION. The *relion_helix_toolbox* program was used to apply real-space symmetry to the final map(49). Finally, a local resolution map was generated using a B-factor of -43 as determined in the automated post-processing job.

A similar processing scheme was followed for the EX-6 ligand-fibril complex. After 3D classification, a final particle set comprised of 64,493 particles was selected for 3D auto-refinement. Two rounds of CTF refinement and Bayesian polishing were performed as for the apo data set. During the final 3D refinement job, helical parameters with a rise of ~4.8 Å, ~9.6 Å, ~14.4 Å, and ~19.2 Å were tested as they represent 1X, 2X, 3X, and 4X helical rise repeats, i.e., fibril layers. Since the position of the ligand was unknown, these additional rise values were tested to resolve ligand binding in either the diagonal, horizontal, or vertical direction, while also testing for possible conformational differences. The final EX-6 ligand-fibril complex map resolved to 2.59 Å with a helical twist of -2.28° and a rise of 9.65 Å. Sharpening was performed in RELION's post-processing scheme with a B-factor of -10. The post-processed map was subjected to real-space symmetrization using the *relion_helix_toolbox* program. Finally, a local resolution map was generated using a B-factor of -10 as applied in the post-processing step (**Figure S4**).

### FRET: Protein Expression, Labeling, and Purification

*Materials*

*E. coli* BL21(DE3) cells were purchased from Stratagene (La Jolla, CA, USA). Milli-Q filtered (18 MΩ) water was used for all solutions (Millipore; Billerica, MA, USA). Cytiva Vivaspin centrifugal filter units (3 kDa MWCO) were purchased from Cytiva Life Sciences. Acridonylalanine (Acd) was synthesized as previously described. (3, 4). BODIPY-FL-maleimide (Bdp-Mal) was purchased from Lumiprobe Life Science Solutions (Hunt Valley, MD, USA). All other reagents and solvents were purchased from Fisher Scientific (Pittsburgh, PA, USA) or Sigma-Aldrich unless otherwise specified. DNA sequencing was performed at the University of Pennsylvania DNA sequencing facility.

*Instruments*

Matrix-assisted laser desorption/ionization with time-of-flight detector (MALDI) mass spectra were acquired on a Bruker rapifleX LRF instrument (Billerica, MA, USA). Fluorescence spectra and protein quantification measurements were collected with a Tecan SPARK plate reader (Mannedorf, Switzerland).

*Construction of pTXB1_αSyn-$C_9$, $TAG_{94}$, $TAG_{114}$, and $TAG_{125}$ plasmids*

Site-directed mutagenesis was performed on three previously constructed pTXB1_αSyn-Mxe-His6 mutants, $TAG_{94}$, $TAG_{114}$, and $TAG_{125}$, containing a TAG mutation for unnatural amino acid incorporation at the indicated site. (4, 5). Quikchange polymerase chain reaction (PCR) using Q5 Hotstart High-Fidelity DNA Polymerase was performed on each TAG-containing mutant to generate the double mutants, and once on the wildtype (αSyn-WT) to develop the single $Ser_9Cys$ mutation, all bearing a polyhistidine-tagged GyrA intein from *Mycobacterium xenopi* (Mxe) fused to the C-terminus of the αSyn to aid in purification.

*αSyn-WT and αSyn-$C_9$ expression*

αS mutant plasmids (pTXB1_αSyn-WT-Mxe-$His_6$ or pTXB1_αSyn-$C_9$-Mxe-$His_6$) were transformed into competent *E. coli* BL21(DE3) cells and plated onto LB agar plates supplemented with Ampicillin (Amp) overnight at 37 °C. Single colonies were used to inoculate 5 mL of LB media supplemented with Amp (100 µg/mL). Primary cultures were incubated at 37 °C with shaking at 250 rpm overnight. A single primary culture was used to inoculate 1 L of LB medium supplemented with AMP (100 µg/mL) and grown at 37 °C with shaking at 250 rpm until the $OD_{600}$ was measured between 0.6 and 0.9. Expression was then induced with IPTG (1 mM), and the cells were placed at 18 °C with shaking at 250 rpm overnight.

*αSyn-$Acd_{94}$, αSyn-$Acd_{114}$, αSyn-$Acd_{125}$, αSyn-$C_9Acd_{94}$, αSyn-$C_9Acd_{114}$ and αSyn-$C_9Acd_{125}$ expression*

αS mutant plasmids (pTXB1_αS-$TAG_{94}$-Mxe-$His_6$, pTXB1_αS-$TAG_{114}$-Mxe-$His_6$, pTXB1_αS-$TAG_{125}$-Mxe-$His_6$, pTXB1_αS-$C_9TAG_{94}$-Mxe-$His_6$, pTXB1_αS-$C_9TAG_{114}$-Mxe-$His_6$, or pTXB1_αS-$C_9TAG_{125}$-Mxe-$His_6$) and an AcdRS/tRNA plasmid (pDule2_MjAcdRSA9) were transformed into competent *E. coli* BL21(DE3) cells and plated onto LB agar plates supplemented with Amp and Streptomycin (Strep), overnight at 37 °C. Single colonies were selected and used to inoculate 5 mL of LB supplemented with ampicillin (Amp) and streptomycin (Strep) (100 µg/mL of each), and the culture was left to shake at 250 rpm and 37 °C overnight. Single primary cultures were used to inoculate 1 L of M9 media (100 mL 10x M9 salts, 2 mM $MgSO_4$, 15 µg/mL $FeCl_2$, 15 µg/mL $ZnCl_2$, 10 nM $CaCl_2$, 0.02% yeast extract, 0.5% glucose) supplemented with Amp and Strep at 37 °C and shaking at 250 rpm until $OD_{600}$ was between 0.6 – 0.9. At this time, Acd amino acid was added (0.5 mM) and the culture was allowed to incubate for 5 more minutes. Expression was then induced by adding IPTG (1 mM), and the culture was incubated at 18 °C with shaking at 250 rpm overnight.

*Purification of αSyn constructs*

Cells were harvested by centrifugation at 4,000 rpm in a GS3 rotor and Sorvall RC-5 centrifuge for 20 minutes at 4 °C. The supernatant was discarded, and the cell pellet was resuspended in 20 mL of Resuspension Buffer (40 mM Tris, pH 8.3) containing PMSF (0.1 mM) and two Roche protease inhibitor cocktail pills (cOmplete™, Mini, EDTA-free Protease Inhibitor Cocktail, glass vial, Roche Cat. #11836170001). Resuspended cells were then lysed by sonication on ice (Amp: 30, Process Time: 5 min, Pulse On: 1 sec, Pulse Off: 2 sec) and then pelleted at 14,000 rpm in a Sorvall RC-5 centrifuge with an SS-34 rotor for 20 minutes at 4 °C. The supernatant was collected and incubated with $Ni^{2+}$-NTA resin (5 mL column volume) for 1 hour at 4 °C with rotation. The slurry was then added to a fritted column and the liquid allowed to flow through. The resin was then washed with 2 x 13 mL of Wash Buffer A (50 mM HEPES, pH

7.5) and 2 x 13 mL of Wash Buffer B (50 mM HEPES, 5 mM imidazole, pH 7.5). The protein constructs were eluted from the resin with 2 x 6 mL of elution buffer (50 mM HEPES, 300 mM imidazole, pH 7.5). The elution fractions were then pooled and treated with β-mercaptoethanol (βME, 200 mM) for intein cleavage at RT for 18 hours on a rotisserie. The resulting cleavage solution was then dialyzed against 20 mM Tris, pH 8.0, overnight. The dialyzed solution was then reapplied to 5 mL of $Ni^{2+}$-NTA and incubated at 4 °C with rotation for 1 hour. The slurry was then applied to a fritted column and allowed to flow through. The resin was washed once with 13 mL of Wash Buffer A and then pooled with the previously collected flowthrough. It was dialyzed against a 20 mM Tris, pH 8.0, buffer at 4 °C overnight. Before FPLC purification, all Cys-containing constructs were treated with TCEP Bond Breaker™ (1 mM) to reduce any disulfides formed between αSyn proteins and/or βME. Each construct was purified by ion exchange chromatography using a HiTrap Q HP column (5 mL) on an ÄKTA FPLC using a 65-minute NaCl gradient (0 to 450 mM NaCl in 20 mM Tris, pH 8.0). The fractions containing pure product were identified using MALDI MS. The αSyn Cys containing mutant constructs were dialyzed against a 20 mM Tris, pH 8.0 buffer at 4 °C overnight. αSyn-WT and αSyn Acd-containing mutant constructs were dialyzed against 1x PBS buffer (137 mM NaCl, 2.7 mM KCl, 10 mM $Na_2HPO_4$, 1.8 mM $KH_2PO_4$, pH 7.4) at 4 °C and further purified via HPLC with a protein C4 column (Phenomenex Jupiter #00G-4168-N0), with pure fractions identified by MALDI MS. Pure protein fractions were pooled and dialyzed against 1x PBS buffer at 4 °C overnight. Following dialysis, proteins were concentrated via centrifugation with a 3 kDa cutoff filter, stored at -80 °C in 1 mL aliquots, and thawed once for experiments. The Cys-containing mutants were immediately subject to labeling following purification.

*αSyn labeling with Bdp-FL-Mal*

Following post-FPLC purification dialysis, the Cys-containing protein solutions (ca. 10-20 mL) were separated into 5 mL aliquots and treated with TCEP Bond Breaker™ (1 mM). To each protein solution, 50 mM Bdp-Mal in DMSO was added to a final concentration of 200 μM dye. The labeling reaction was allowed to proceed for 3 hours on a rotisserie at RT or overnight at 4 °C. Each step was monitored by MALDI MS. After labeling was deemed complete, each solution was dialyzed against a 1x PBS, pH 7.4 at 4 °C for 4 hours to remove excess dye. The dialyzed solution was concentrated via centrifugation with a 3 kDa cutoff filter (Cytiva # 28932358). The concentrated solution was then additionally purified via HPLC with a protein C4 column (Phenomenex Jupiter #00G-4168-N0) and pure protein fractions were identified via MALDI MS. Pure protein fractions were pooled and dialyzed against 1x PBS buffer at 4 °C overnight. Following dialysis, proteins were concentrated via centrifugation with a 3 kDa cutoff filter, stored at -80 °C in 1 mL aliquots, and thawed once for FRET experiments. MALDI MS confirmation of all WT and labeled protein identities is given in **Table S2**.

**Table S2** Protein MALDI Masses.

| Protein | Calc. $[M+H]^+$ | Obs. $[M+H]^+$ |
|---|---|---|
| αS-WT | 14460 | 14459 |
| αS-$C_9$ | 14476 | 14481 |
| αS-$C^{Bdp}{}_9$ | 14890 | 14890 |
| αS-$Acd_{94}$ | 14578 | 14579 |
| αS-$Acd_{114}$ | 14595 | 14598 |
| αS-$Acd_{125}$ | 14561 | 14566 |
| αS-$C_9Acd_{94}$ | 14593 | 14592 |
| αS-$C^{Bdp}{}_9Acd_{94}$ | 15008 | 15005 |
| αS-$C_9Acd_{114}$ | 14611 | 14616 |
| αS-$C^{Bdp}{}_9Acd_{114}$ | 15026 | 15023 |
| αS-$C_9Acd_{125}$ | 14577 | 14575 |
| αS-$C^{Bdp}{}_9Acd_{125}$ | 14992 | 14990 |

**FRET: Spectra Measurements and Analysis**

*Fibril preparation*
Protein construct concentrations were determined by DC assay analysis. αSyn monomer was combined to a final concentration of 100 μM with 95:5 WT to labeled construct (or 100 μM pure WT for control experiments). Each aliquot was treated with sodium azide to a final concentration of 0.2% to restrict bacterial growth. The tubes were sealed with Teflon and Parafilm and incubated at 37 °C with shaking at 1300 rpm for 5 days. The fibrils were then pelleted, and the supernatant was removed to eliminate any unincorporated monomer. Pellets were then reconstituted in fresh 1x PBS with 0.2% sodium azide.

*Fluorescence measurements*
αSyn fibrils were diluted to a final concentration of 10 μM (relative to αS monomer) into 1x PBS buffer and pipetted in triplicate into nonsterile Greiner black, flat μClear, 96 well half area microplates (#675096) to a final volume of 100 μL. Additionally, Bdp dye, Acd amino acid, αSyn-$C^{Bdp}{}_9$, αSyn-$δ_{94}$, αSyn-$δ_{114}$, αSyn-$δ_{125}$, αSyn-$C^{Bdp}{}_9δ_{114}$ and αSyn-$C^{Bdp}{}_9δ_{125}$ monomers were all diluted to 0.5 μM and subjected to the same conditions as below to act as controls and as comparison for data analysis. Before the addition of small molecules, fluorescence intensity values were obtained for each construct to optimize gain and Z-position using $λ_{ex}$ = 385 ± 5 nm and a reading at $λ_{em}$ = 420 ± 5 nm, as Acd is expected to have the highest fluorescence intensity. Additionally, fluorescence intensity was measured, exciting at 490 ± 5 nm and reading at 515 ± 5 nm to optimize gain and Z-position for direct Bdp excitation. T 0 h emission scans were obtained for each construct before small molecule addition using the parameters previously acquired and optimized, exciting at $λ_{ex}$ = 410 ± 5 nm, scanning from $λ_{em}$ = 425 to 600 ± 5 nm, in 1 nm increments, and a 40 μs integration time. Direct excitation of Bdp was also acquired at $λ_{ex}$ = 490 ± 5 nm and measured from

$\lambda_{em}$ = 505 to 600 ± 5 nm using the same step size and integration time as previously mentioned. Small molecule was then added in triplicate to a final concentration of 10 μM. The plates were then covered with an anti-evaporation guard and placed at 37 °C with shaking at 500 rpm. Measurements were taken using the same parameters at 4 hours, and 24 hours after small molecule addition.

*FRET spectra analysis*

Förster resonance energy transfer (FRET) data were fit as previously described (5, 6). Triplicates of raw spectra were averaged and background corrected by subtraction of the corresponding averaged spectra of WT fibrils. Inspection of corrected spectra shows quenching of Acd and an increase in the Bdp signal in the doubly labeled (DL) spectra vs the sum of the Acd-only and Bdp-only control spectra, indicating FRET (**Figure S8**).To determine FRET efficiency, or $E_{FRET}$, the linear combination of the Acd or Bdp singly labeled spectra was fit to the acquired DL spectra by adjusting A and B coefficients to minimize their square difference, as seen in Equation 1 below, in which $I(\lambda)_{DL}$, $I(\lambda)_{Acd}$, and $I(\lambda)_{Bdp}$ represent the fluorescence intensity of the DL fibrils, the Acd-only, and the Bdp-only, respectively at a given wavelength. Fitted spectra are shown in **Figure S9**.

$$\sum_{\lambda}(I(\lambda)_{DL} - (AI(\lambda)_{Acd} + BI(\lambda)_{Bdp}))^2 \quad \text{(Eq. 1)}$$

$E_{FRET}$ was then converted to distance $R$ using Equation 2 below, where $R$ is the distance between the fluorophores, and $R_0$ is the FRET distance of the Bdp-Acd fluorophore pair, as determined by Equation 3.

$$E_{FRET} = \frac{1}{1+(\frac{R}{R_0})^6} \quad \text{(Eq. 2)}$$

$$R_0^6 = \frac{9000(\ln 10)\kappa^2\Phi_D J}{128\pi^5 n^4 N_A} \quad \text{(Eq. 3)}$$

In Equation 3, $\kappa^2$ represents the orientation parameter and is assumed to be 2/3, $\Phi_D$ is the quantum yield of the donor fluorophore in its respective protein environment, determined by comparing to emission of the free Acd amino acid in water (Φ = 0.95), $J$ represents the overlap integral of the emission of Acd and the absorbance of Bdp, determined to be $8.18 \times 10^{14}$ $M^{-1}{*}cm^{-1}{*}nm^4$, $n$ is the refractive index of water 1.33, and $N_A$ is Avogadro's number. Quantum yield values and their respective $R_0$ values for each Acd-only containing construct are listed in **Table S3**. $E_{FRET}$ and corresponding $R$ values for each FRET pair can be found in **Table S4**.

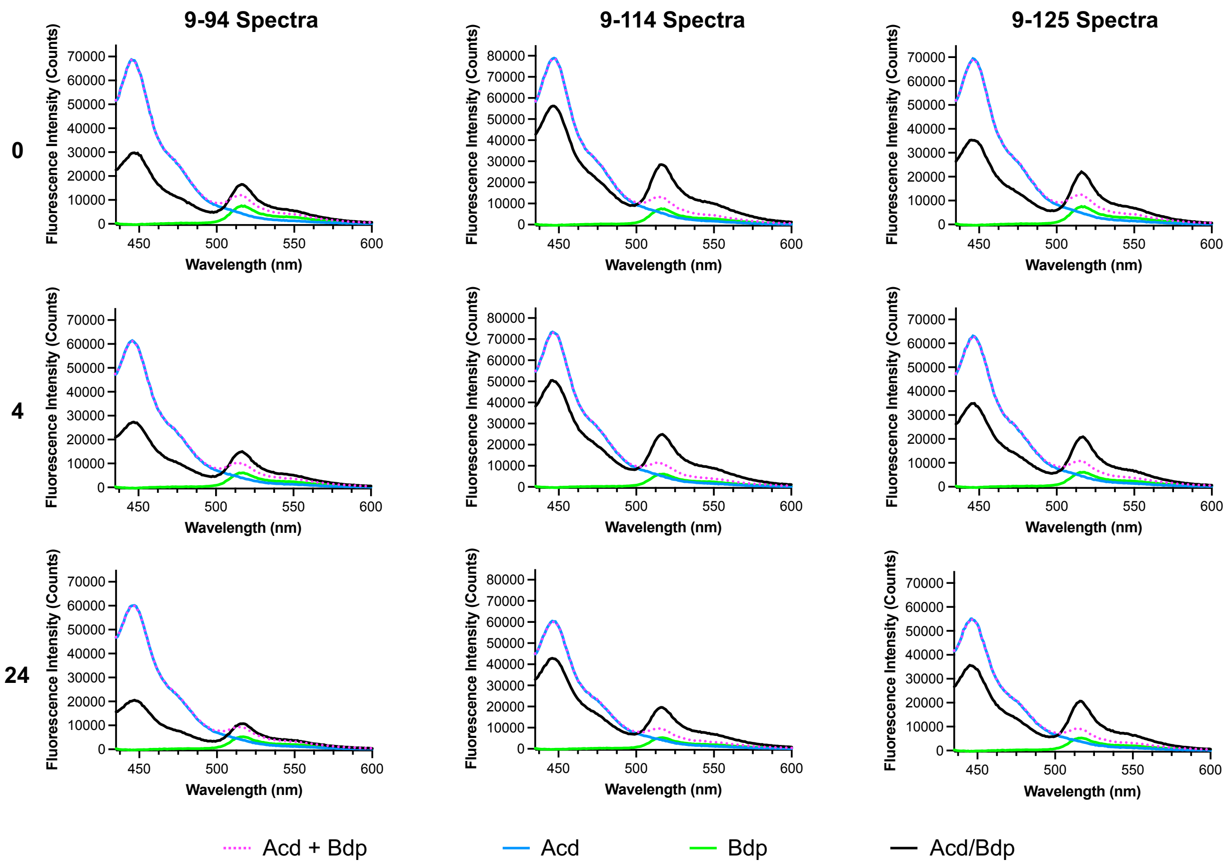


**Figure S7** αSyn Fibril Spectra. Donor-only and acceptor-only spectra as well as their sum compared to the double-labeled spectrum for untreated (0) fibrils and fibrils treated with EX-6 for 4 hours or 24 hours. Donor-acceptor pairs: $Acd_{94}$-$Bdp_9$, $Acd_{114}$-$Bdp_9$, $Acd_{125}$-$Bdp_9$.

**Table S3** Donor quantum yields and Förster radii[a]

| Condition | 94 | 114 | 125 |
|---|---|---|---|
| Untreated $\Phi_D/R_0$ (Å) | 0.65/46 | 0.77/48 | 0.67/47 |
| 4 h $\Phi_D/R_0$ (Å) | 0.59/45 | 0.71/47 | 0.66/47 |
| 24 h $\Phi_D/R_0$ (Å) | 0.58/45 | 0.59/45 | 0.61/46 |

[a]Estimated variance: $\Phi_D$ ± 0.03, $R_0$ ± 1 Å.

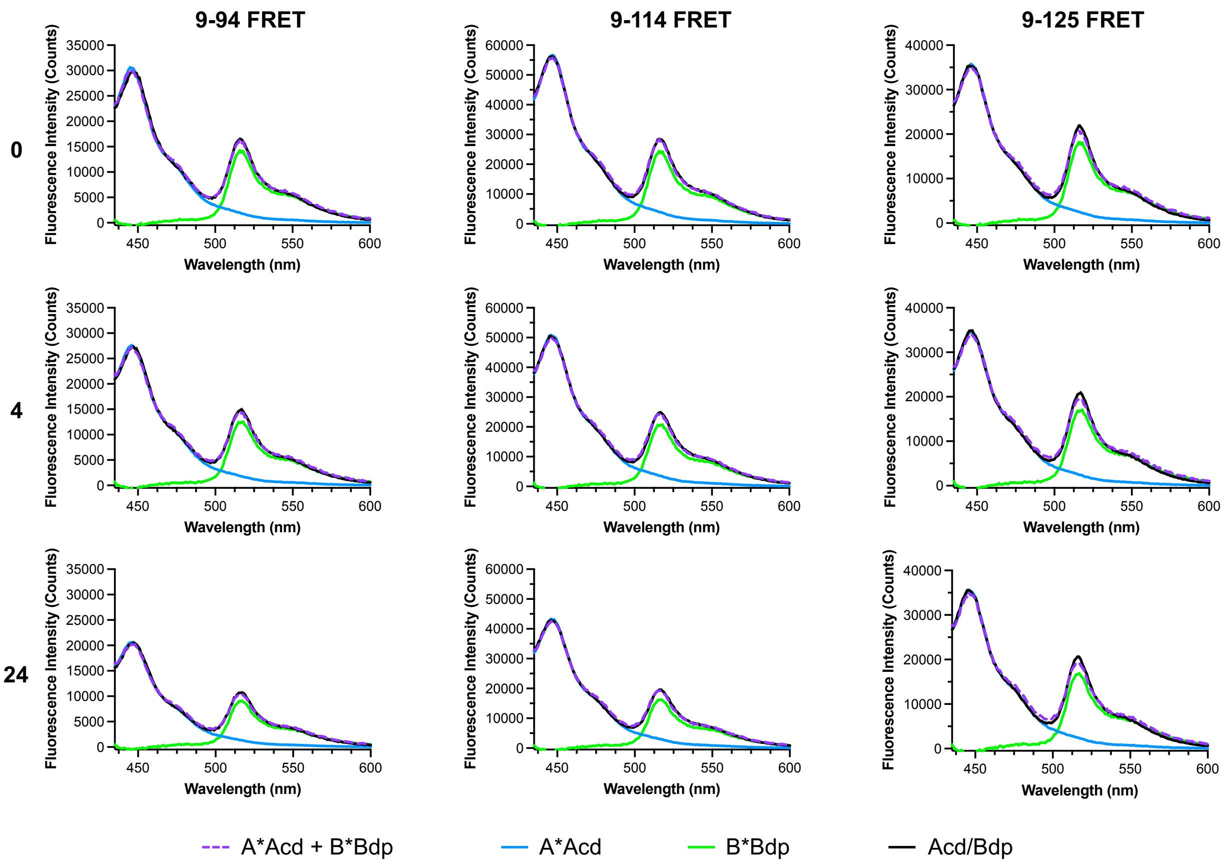


**Figure S8** αSyn Fibril FRET. $E_{\mathrm{FRET}}$ calculations for fibril data showing the weighted donor-only and acceptor-only spectra as well as their sum compared to the double-labeled spectrum for untreated (0) fibrils and fibrils treated with EX-6 for 4 hours or 24 hours. Donor-acceptor pairs: $\mathrm{Acd}_{94}$-$\mathrm{Bdp}_{9}$, $\mathrm{Acd}_{114}$-$\mathrm{Bdp}_{9}$, $\mathrm{Acd}_{125}$-$\mathrm{Bdp}_{9}$.

**Table S4** FRET efficiencies and distance calculations[a]

| Condition | 9-94 | 9-114 | 9-125 |
|---|---|---|---|
| Untreated $E_{\mathrm{FRET}}$/R (Å) | 0.55/45 | 0.28/56 | 0.49/47 |
| 4 h $E_{\mathrm{FRET}}$/R (Å) | 0.55/44 | 0.31/54 | 0.45/49 |
| 24 h $E_{\mathrm{FRET}}$/R (Å) | 0.66/41 | 0.28/53 | 0.36/50 |

[a]Estimated variance: $E_{\mathrm{FRET}}$ measurements ±0.05. Distance calculations ±2 Å.

**Photo-crosslinking: Probe Synthesis**

*Chemical reagents and instruments*

Chemicals were obtained from commercial sources and used without further purification. Solvents were purchased from commercial sources and used as received unless stated otherwise. Reactions were performed at room temperature unless stated otherwise. Reactions were monitored by thin-layer chromatography (TLC) on pre-coated silica 60 F254 aluminum plates (MilliporeSigma, Burlington, MA, USA), and spots were visualized by ultraviolet (UV) light. Evaporation of solvents was performed under reduced pressure at 40 °C using a rotary evaporator. Flash column chromatography was performed on a Biotage® (Charlotte, NC, USA) Isolera One system equipped with Biotage® SNAP KP-Sil cartridges. Nuclear magnetic resonance (NMR) spectroscopy was performed on a Bruker (Billerica, MA, USA) Avance Neo 600 (600 MHz for $^{1}H$ and 150 MHz for $^{13}C$) with chemical shifts (δ) reported in parts per million (ppm) relative to the solvent ($CDCl_3$, $^{1}H$ 7.26 ppm, $^{13}C$ 77.16 ppm; dimethyl sulfoxide (DMSO)-$d_6$, $^{1}H$ 2.50 ppm, $^{13}C$ 39.52 ppm). Low resolution liquid chromatography mass spectrometry (LCMS) was carried out using a Waters (Milford, MA, USA) SQD equipped with an Acquity UPLC instrument in positive-ion mode. High resolution mass spectrometry (HRMS) for small molecules was obtained on a Waters LCT Premier XE LC/MS system or Bruker Sci-Max. Python version 3.10.13 was used for data analysis scripts. ChatGPT drafted visualization scripts, which were thoroughly checked and edited for implementation details.

*Synthesis of precursor 1 (**CLX-P-1**)*

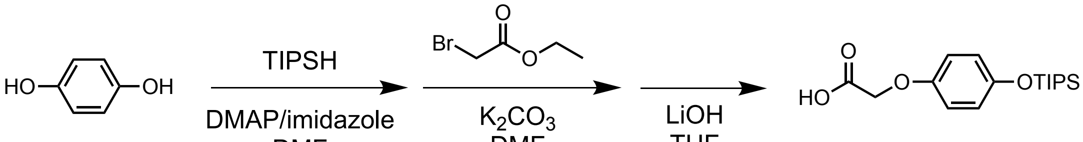


**CLX-P1**

To a flame dried round bottom flask, hydroxyquinone (90.8 mmol, 2 equiv.), imidazole (136.0 mmol, 3 equiv.), and 4-dimethylaminopyridine (4.5 mmol, 0.05 equiv.), were added. The flask was charged with argon and DMF (200 mL) was added to the flask with strong stirring. Triisopropylsilyl chloride (45 mmol, 1 equiv.) was added dropwise to the flask over 5 minutes. The reaction was left at room temperature for 16 hours. The reaction mixture was combined with saturated $Na_2CO_3$ in water (100 mL) and extracted with EtOAc (3x 50 mL). The combined organic phase was washed with water (2x 30 mL), saturated brine (30 mL), dried over $Na_2SO_4$, and dried under reduced pressure. The crude mixture was subjected to flash chromatography with a 15:90 EtOAc:Hexane (v/v) gradient to obtain 4-((triisopropylsilyl)oxy)phenol.

To a flame-dried round-bottom flask, 4-((triisopropylsilyl)oxy)phenol (26.4 mmol, 1 equiv.) and anhydrous $K_2CO_3$ (79.3 mmol, 3 equiv.) were added. The flask was evacuated and backfilled with argon three times, then anhydrous DMF (60 mL) was introduced with vigorous stirring. The mixture was warmed to 70 °C, and ethyl bromoacetate (79.3 mmol, 3 equiv.) was added drop-wise over 10 minutes. To the reaction mixture, 50 mL of water was added and extracted with EtOAc (3x 30 mL). The combined organic layers were washed with water (2 × 20 mL) and saturated brine (20 mL), dried over $Na_2SO_4$, filtered, and concentrated under reduced pressure. The crude residue was purified by flash chromatography with 10:90 EtOAc:Hexanes (v/v) to obtain methyl 2-(4-((triisopropylsilyl)oxy)phenoxy)acetate.

Methyl 2-(4-((triisopropylsilyl)oxy)phenoxy)acetate (19.2 mmol, 1 equiv.) was added to a flask open to air with THF (30 mL) under strong stirring. In a separate beaker, water (30 mL) was mixed with LiOH (96.1 mmol, 5 equiv.) and added dropwise to the organic solution over 10 minutes. The reaction was allowed to stir at room temperature for 2 hours. The THF was removed under reduced pressure. The pH of the solution was then adjusted to 2.0 using 1M HCl in water and the product then precipitated from the solution. The solution was extracted with EtOAc (3x 30 mL) and the combined organic layers were washed with brine containing 0.1 HCl. The organic solution was dried over $Na_2SO_4$ and concentrated under reduced pressure. The crude mixture was subjected to flash chromatography utilizing 5:95 MeOH:DCM (v/v) to afford **CLX-P-1.**

**CLX-P-1** Characterization: White solid; TLC (MeOH:DCM 5:95 v/v) $R_f$: 0.42; 62% yield (over 3 steps). $^{1}H$ NMR (600 MHz, $CDCl_3$) δ 6.84 – 6.77 (m, 4H), 4.61 (s, 2H), 1.23 (dt, *J* = 14.9, 7.5 Hz, 3H), 1.09 (d, *J* = 7.4

Hz, 18H). $^{13}$C NMR (150 MHz, DMSO) δ 170.82, 152.56, 149.87, 120.48, 115.78, 65.44, 18.21, 12.46. HRMS (*m/z*): calculated for $C_{17}H_{29}O_4Si$ [M+H]$^+$ 325.1830 found 325.1830

*Synthesis of precursor 2 (**CLX-P-2**)*

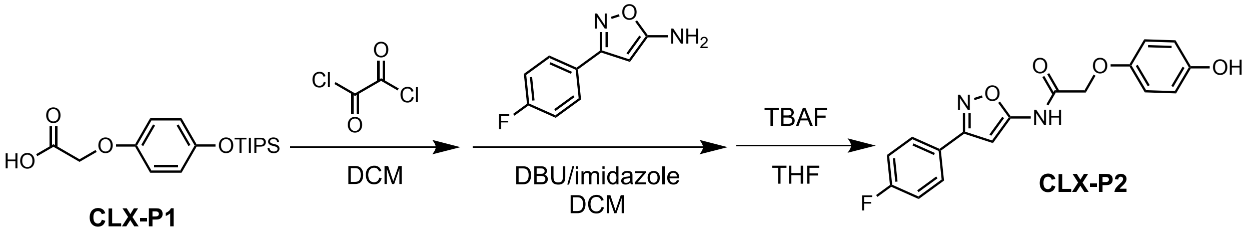


To a flame dried round bottom flask equipped with a stir bar, **CLX-P-1** (1.66 mmol, 5 equiv.) was added. The flask was backfilled with argon 3x. DCM (20 mL) was added to the flask and the solution was cooled to 0 °C by an ice water bath. Oxalyl chloride (2.30 mmol, 7.5 equiv.) was added dropwise over 5 minutes. The solution was left to stir for 2 hours, after which, the solvent was evaporated, and the reaction flask was placed under argon. In a separate vial 1,8-Diazabicyclo[5.4.0]undec-7-ene (DBU, 3.32 mmol, 10 equiv.) and imidazole (1.66 mmol, 5 equiv.) were mixed in DCM (20 mL) and added to the reaction flask. 3-(4-fluorophenyl)isoxazol-5-amine (0.33 mmol, 1 eq) was added to the flask over 5 minutes and the reaction was left to stir for 2 hours. $NaHCO_3$ (20 mL) was added to the solution and the organic layer was washed with DCM (3x 10 mL). The organic layer was dried over $Na_2SO_4$ and the solvent was evaporated under reduced pressure. The crude mixture was subjected flash chromotagraphy (30:70 EtOAc:Hexanes v/v), however, two compounds co-eluted and was transferred as a mixture to the next step. The flask containing the protected material was sparged with argon (3x). To the flask, THF (15 mL) was added and the reaction mixture was cooled to 0 °C over a water bath. Tetra-n-butylammonium fluoride (0.66 mmol, 2 equiv.) was added dropwise over one minute. The reaction was stirred continuously for 1 hour. The solvent was evaporated and the crude mixture was subjected to flash chromatography (60:40 EtOAc:Hexanes v/v) to obtain **CLX-P-2**.
**CLX-P-2** Characterization: Off-White Solid; TLC (50:50 EtOAc/Hexanes (v/v) $R_f$: 0.34; 55% yield (over 2 steps). $^1$H NMR (600 MHz, DMSO) δ 11.92 (s, 1H), 9.01 (s, 1H), 7.96 – 7.91 (m, 2H), 7.34 (t, *J* = 8.8 Hz, 2H), 6.87 – 6.79 (m, 3H), 6.73 – 6.67 (m, 2H), 4.71 (s, 2H). $^{13}$C NMR (150 MHz, DMSO) δ 166.56, 164.50, 162.87, 162.20 (d, *J* = 3.6 Hz), 152.32, 150.97, 129.36 (d, *J* = 8.6 Hz), 125.72, 116.57 (d, *J* = 21.8 Hz), 116.25 – 116.11 (m), 87.09, 67.82. HRMS (*m/z*): calculated for $C_{17}H_{14}FN_2O_4^+$ [M+H]$^+$ 329.0932 found 329.0940

*Synthesis of **EX-6-CLX***

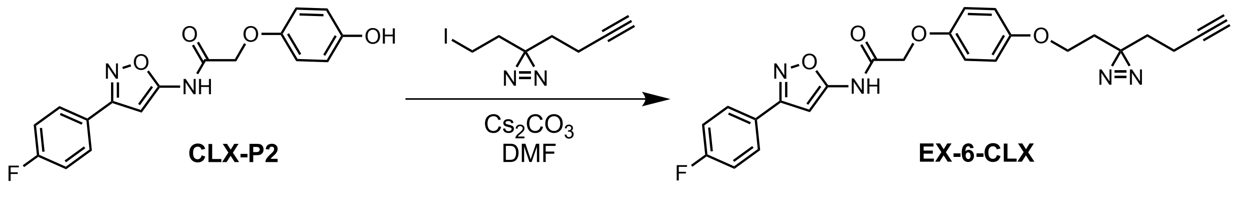


To a flame dried flask containing a stir bar, **CLX-P-2** (60.9 µmol, 1 equiv.) and $Cs_2CO_3$ (365.5 µmol, 4 equiv.) was added to the flask. The flask was purged of air under vacuum and backfilled with argon (3x). DMF (5 mL) was added to the flask and strong stirring was enabled. 3-(but-3-yn-1-yl)-3-(2-iodoethyl)-3H-diazirine (243.7 µmol, 4 equiv.) was added dropwise to the flask over 2 minutes. The reaction was stirred for 16 hours after which EtOAc (10 mL) and water (15 mL) was added. The water layer was washed with EtOAc (10 mL). The combined organic layers were washed once with brine (10 mL) and dried over $Na_2SO_4$. The crude mixture was concentrated under reduced pressure and subjected to flash chromatography (50:50 EtOAc:Hexanes v/v) to obtain **EX-6-CLX**.
**EX-6-CLX** Characterization: Yellow solid; TLC (30:70 EtOAc:Hexanes v/v) $R_f$ 0.29; 65% yield. $^1$H NMR (600 MHz, $CDCl_3$) δ 9.17 (s, 1H), 7.85 – 7.79 (m, 2H), 7.20 – 7.12 (m, 2H), 6.97 – 6.85 (m, 4H), 6.79 (s, 1H), 4.65 (s, 2H), 3.79 (t, *J* = 6.2 Hz, 2H), 2.07 (td, *J* = 7.5, 2.7 Hz, 2H), 1.99 (t, *J* = 2.7 Hz, 1H), 1.89 (t, *J* = 6.2 Hz, 2H), 1.74 (t, *J* = 7.5 Hz, 2H). $^{13}$C NMR (150 MHz, $CDCl_3$) δ 164.95, 163.05, 159.77, 154.30, 151.10,

128.92 (d, $J$ = 8.7 Hz), 125.27 (d, $J$ = 3.2 Hz), 116.20 (d, $J$ = 21.8 Hz), 116.11, 116.04, 87.61, 82.88, 69.35, 68.09, 63.31, 33.06, 32.84, 26.81, 18.05, 13.45. HRMS (*m/z*): calculated for 449.1625 $[M+H]^+$ found 449.1613.

**Photo-crosslinking: MS Analysis**

*General Information*
Buffers were made with MilliQ filtered (18 MΩ) water (Millipore; Billerica, MA, USA). DC protein assay kits were purchased from Bio-Rad (Bio-Rad; Hercules, CA, USA). DC assay protein quantitation was conducted using a Tecan Spark microplate reader (Tecan; Mannedorf, Switzerland). Matrix-assisted laser desorption/ionization (MALDI) mass spectrometry (MS) data were collected with a Bruker Rapiflex MALDI-TOF/TOF mass spectrometer (Billerica, MA, USA).

*Crosslinking*
EX-6-CLX was prepared in a DMSO stock solution to a final concentration of 10 mM and was diluted to the appropriate concentrations using DMSO. All experiments involving EX-6-CLX had a final concentration of 1% DMSO. See **Table S5** for the conditions of each experiment. For a detailed fibril sample preparation procedure, please refer to the main text. **Figure S10, Figure S11, and Figure S12** show representative MS data for crosslinking under conditions 1, 3, and 4, respectively. Representative MS data for condition 2 are shown In the main text. **Figure S13** shows two accepted mechanisms for diazirine crosslinking with associated amino acid selectivity (7, 8). We have primarily observed crosslinking at Asp and Glu residues, as seen in MS2 spectra shown in **Figures S11 - S14** show MS2 of tryptic peptides which are crosslinked to EX-6-CLX. A complete list of MS peaks observed under condition 2 is given in **Table S6**. A complete list of MS peaks observed under condition 1 is given in **Table S7.** Peaks were assigned using fragments from a theoretical trypsin digest of the human αSyn sequence using the Protein Prospector website with up to two missed cleavages and oxidation modifications turned on. Additional candidate m/z values were generated by using Protein Prospector to predict isotopomer distributions and considering sodium adducts, crosslinking of EX-6-CLX (+420.1485), and EX-6-CLX fragments (9). All MS spectra were acquired in positive ion mode with 5000 shots at 45% laser power. A signal to noise of 6 was implemented along with a ppm difference between matching peaks of 50. For all MS2 spectra, 2000 fragment shots of the parent ion were collected and added to 4000 shots of the fragments produced through post-source decay and collision induced dissociation from argon gas.

**Table S5** Crosslinking sample preparation

| Condition Number | Concentration | [EX-6-CLX] (µM) | Protease | Final Volume (µL) |
|---|---|---|---|---|
| 1 | Fibril (10 µM) | 100 | Trypsin/Lys-C | 100 |
| 2 | Fibril (1 µM) | 0.5 | Trypsin/Lys-C | 1000 |
| 3 | Monomer | 100 | Trypsin/Lys-C | 100 |
| 4 | Monomer | 0.5 | Trypsin/Lys-C | 1000 |

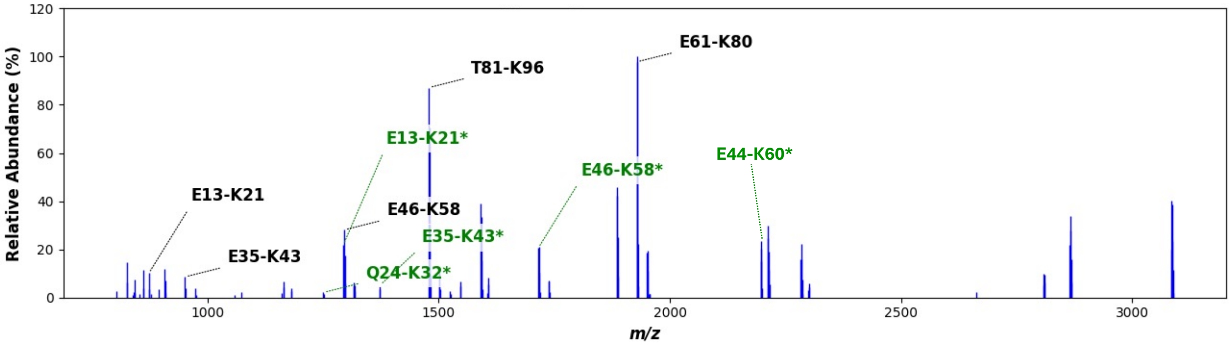


**Figure S9** High concentration of EX6-CLX crosslinking to fibrillar αSyn (Sample 1). Representative MS1 of tryptic peptides resulting from digestion.

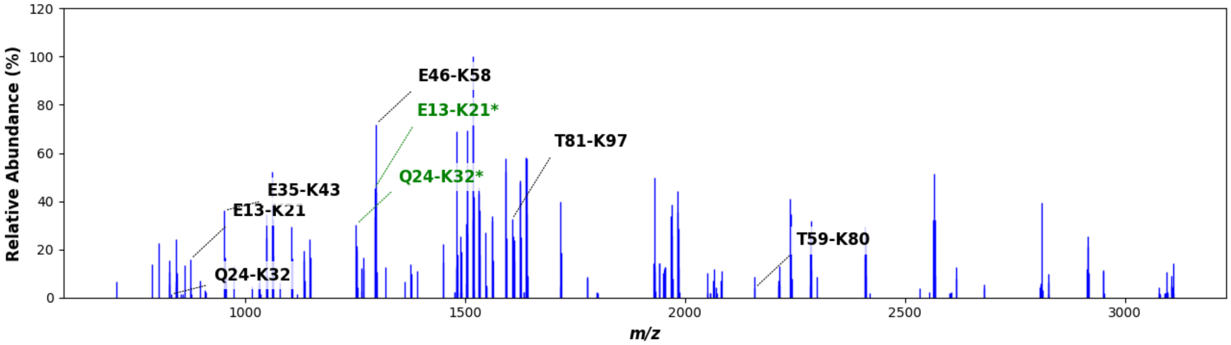


**Figure S10** High concentration of EX6-CLX crosslinking to monomeric αSyn (Sample 3). Representative MS1 of tryptic peptides resulting from digestion.

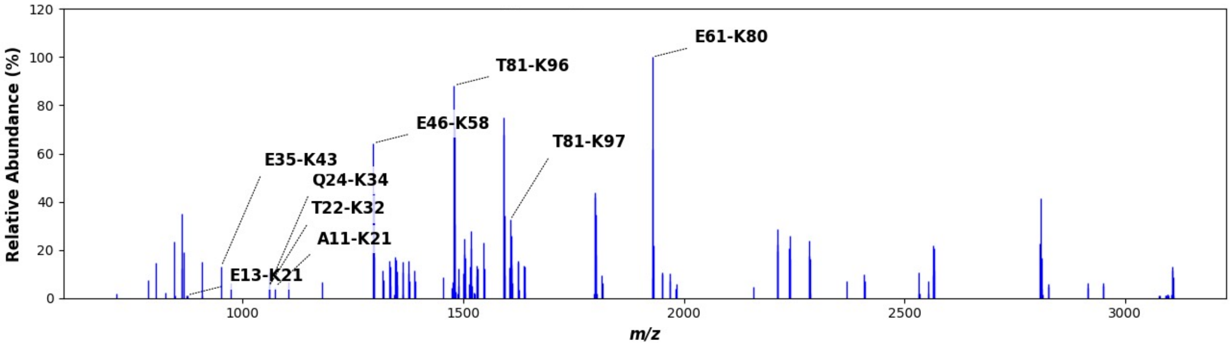


**Figure S11** Low concentration of EX6-CLX crosslinking to monomeric αSyn (Sample 4). Representative MS1 of tryptic peptides resulting from digestion.

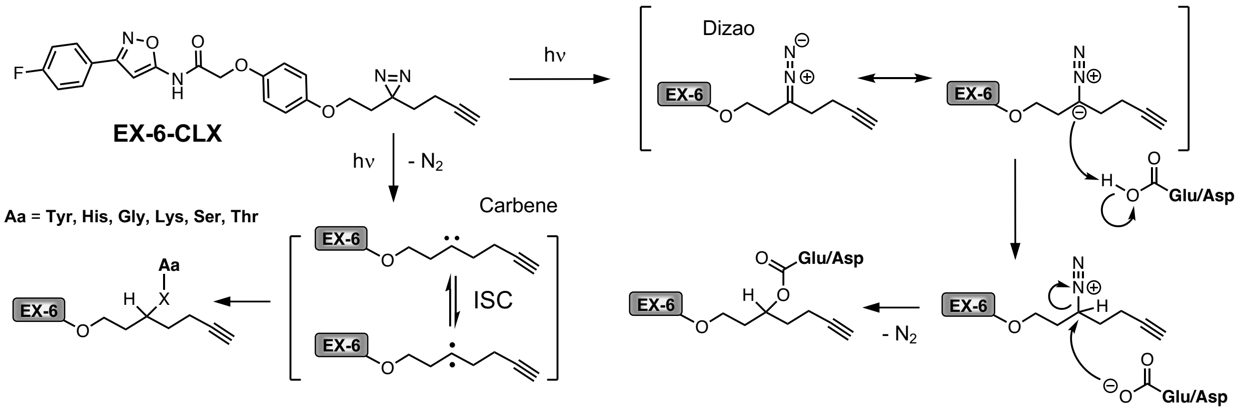


**Figure S12** Crosslinking mechanisms of EX-6-CLX. Diazirine crosslinking can occur through diazo or carbene mechanisms, with different amino acid selectivity.

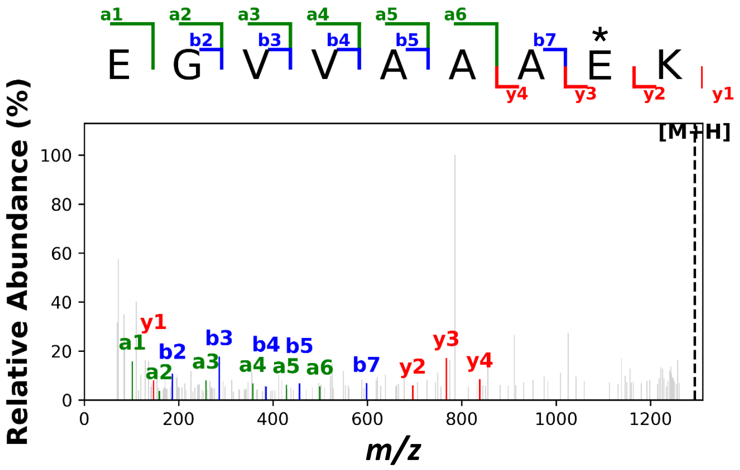


**Figure S13** Representative MS2 of the sequence [13]EGVVAAAE*K[21] from high concentration EX6-CLX crosslinking to fibrillar αSyn. The asterisk indicates the crosslinked residue used in MS2 comparison to a theoretical digest. Ions found, a, b, and y, are shown in green, blue, and red respectively.

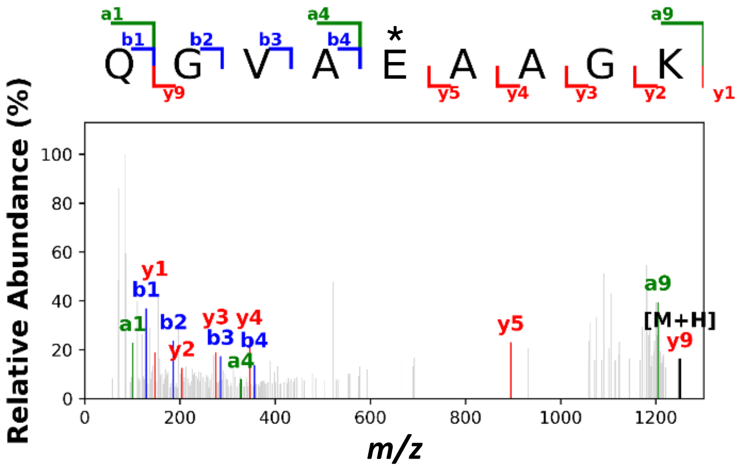


**Figure S14** Representative MS2 of the sequence [24]QGVAE*AAGK[32] from high concentration EX6-CLX crosslinking to fibrillar αSyn. The asterisk indicates the crosslinked residue used in MS2 comparison to a theoretical digest. Ions found, a, b, and y, are shown in green, blue, and red respectively.

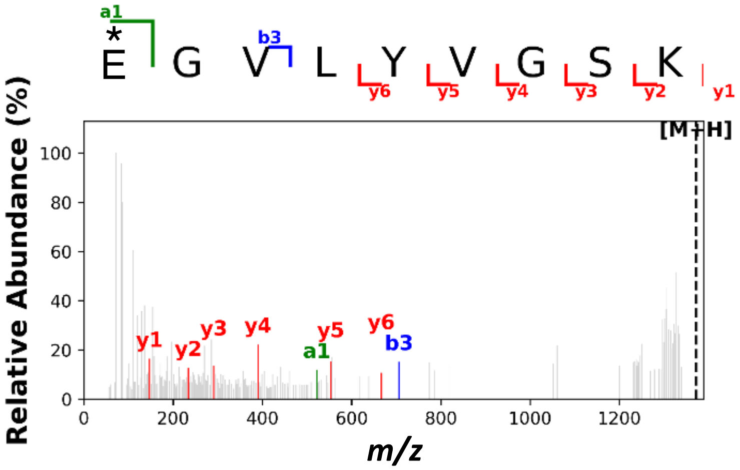


**Figure S15** Representative MS2 of the sequence [35]E*GVLYVGSK[43] from high concentration EX6-CLX crosslinking to fibrillar αSyn. The asterisk indicates the crosslinked residue used in MS2 comparison to a theoretical digest. Ions found, a, b, and y, are shown in green, blue, and red respectively.

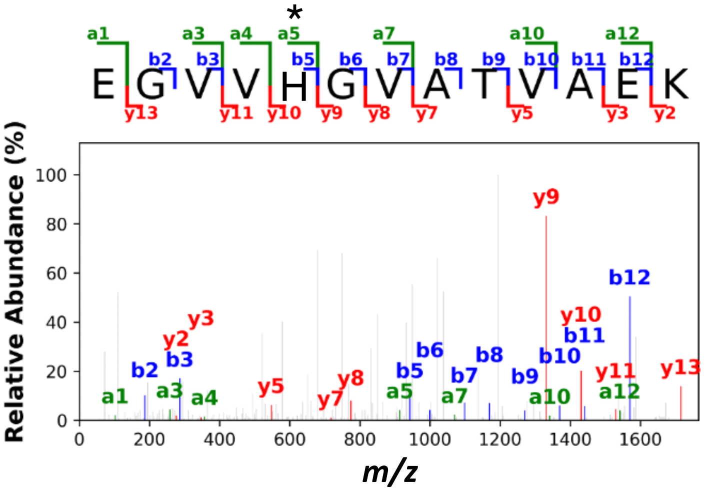


**Figure S16** Representative MS2 of the sequence [46]EGVVH*GVATVAEK[58] from high concentration EX6-CLX crosslinking to fibrillar αSyn. The asterisk indicates the crosslinked residue used in MS2 comparison to a theoretical digest. Ions found, a, b, and y, are shown in green, blue, and red respectively.

**Table S6** Fibril crosslinking with 0.5 μM EX-6-CLX

| Obs m/z | Intensity (Counts) | Calc m/z | Tryptic Peptide[a] | Notes[c] | Difference (ppm) |
|---|---|---|---|---|---|
| 845.099 | 25090 | | | | |
| 861.070 | 246033 | | | DMSO | |
| 862.072 | 106721 | | | DMSO | |
| 863.076 | 11384 | | | | |
| 867.088 | 34331 | | | | |
| 877.042 | 39804 | | | DMSO | |
| 907.345 | 32985 | | | DMSO | |
| *1072.091* | *21614* | *1072.600* | *A11-K21* | *Unmodified* | *474* |
| 1293.612 | 33073 | 1293.616 | E13-K21 | +EX-6-CLX | 3 |
| 1294.616 | 17790 | 1294.619 | E13-K21[b] | +EX-6-CLX | 2 |
| 1295.681 | 62383 | 1295.695 | E46-K58 | Unmodified | 11 |
| 1296.690 | 38402 | 1296.698 | E46-K58[b] | Unmodified | 7 |
| 1478.782 | 220876 | 1478.785 | T81-K96 | Unmodified | 2 |
| 1479.781 | 174893 | 1479.788 | T81-K96[b] | Unmodified | 4 |

[a]All m/z calculated for $[M+H]^+$ species unless otherwise indicated.
[b]Minor isotopomer. Isotopomer m/z determined using Protein Prospector.
[c]DMSO indicates that although the peak was identified it is also observed in DMSO control.

**Table S6 cont'd.** Fibril crosslinking with 0.5 μM EX-6-CLX

| Obs m/z | Intensity (Counts) | Calc m/z | Tryptic Peptide[a] | Notes[c] | Difference (ppm) |
|---|---|---|---|---|---|
| 1480.785 | 72459 | 1480.791 | T81-K96[b] | Unmodified | 4 |
| 1481.788 | 15268 | 1481.793 | T81-K96[b] | Unmodified | 4 |
| 1500.771 | 19196 | 1500.767 | T81-K96+Na | Unmodified | -2 |
| 1501.778 | 10670 | 1501.770 | T81-K96+Na[b] | Unmodified | -5 |
| 1590.940 | 122293 | | | DMSO | |
| 1591.940 | 107776 | | | DMSO | |
| 1592.940 | 51194 | | | DMSO | |
| *1606.924* | *15878* | *1606.880* | *T81-K97* | *Unmodified* | *-28* |
| *1607.928* | *5635* | *1607.883* | *T81-K97[b]* | *Unmodified* | *-28* |
| 1715.840 | 31461 | 1715.844 | E46-K58 | +EX-6-CLX | 2 |
| 1716.847 | 33051 | 1716.838 | E46-K58[b] | +EX-6-CLX | -5 |
| 1717.852 | 13276 | 1717.849 | E46-K58[b] | +EX-6-CLX | -1 |
| 1737.827 | 3443 | 1737.826 | E46-K58[b] | +EX-6-CLX | -1 |
| 1884.830 | 127865 | | | DMSO | |
| 1885.833 | 138320 | | | DMSO | |
| 1886.837 | 70502 | | | DMSO | |
| 1887.836 | 21497 | | | DMSO | |
| 1928.036 | 126038 | 1928.045 | E61-K80 | Unmodified | 4 |
| 1929.039 | 139078 | 1929.048 | E61-K80[b] | Unmodified | 4 |
| 1930.039 | 73458 | 1930.050 | E61-K80[b] | Unmodified | 6 |
| 1931.040 | 23335 | 1931.053 | E61-K80[b] | Unmodified | 7 |
| 1950.023 | 20662 | 1950.027 | E61-K80+Na | Unmodified | 2 |
| 1951.028 | 23396 | 1951.030 | E61-K80+Na[b] | Unmodified | 1 |

[a]All m/z calculated for $[M+H]^+$ species unless otherwise indicated.
[b]Minor isotopomer. Isotopomer m/z determined using Protein Prospector.
[c]DMSO indicates that although the peak was identified it is also observed in DMSO control.

**Table S7** Fibril crosslinking with 100 μM EX-6-CLX

| Obs m/z | Intensity (Counts) | Calc m/z | Tryptic Peptide[a] | Notes[c] | Difference (ppm) |
|---|---|---|---|---|---|
| 825.098 | 214045 | | | | |
| 826.100 | 84539 | | | | |
| 841.074 | 45015 | | | | |
| 842.509 | 13921 | | | DMSO | |
| 861.071 | 64064 | | | DMSO | |
| 862.071 | 10790 | | | DMSO | |
| 873.469 | 27095 | 873.468 | E13-K21 | Unmodified | -1 |
| 907.345 | 35631 | | | DMSO | |
| 951.512 | 17096 | 951.515 | E35-K43 | Unmodified | 3 |
| 1164.534 | 27101 | | | DMSO | |
| 1165.536 | 8862 | | | DMSO | |
| 1293.616 | 79371 | 1293.616 | E13-K21 | Modified | 0 |
| 1294.617 | 55961 | 1294.619 | E13-K21[b] | Modified | 2 |
| 1295.680 | 85241 | 1295.695 | E46-K58 | Unmodified | 12 |
| 1296.693 | 46742 | 1296.698 | E46-K58[b] | Unmodified | 4 |
| 1315.602 | 24765 | 1315.598 | E13-K21 | Modified | -3 |
| 1316.606 | 10939 | 1316.601 | E13-K21[b] | Modified | -4 |
| 1317.650 | 8895 | 1317.604 | E13-K21[b] | Modified | -35 |
| 1371.669 | 6080 | 1371.663 | E35-K43 | Modified | -5 |
| 1478.788 | 302292 | 1478.785 | T81-K96 | Unmodified | -2 |
| 1479.788 | 236384 | 1479.788 | T81-K96[b] | Unmodified | 0 |
| 1480.787 | 98947 | 1480.791 | T81-K96[b] | Unmodified | 2 |
| 1481.792 | 25989 | 1481.793 | T81-K96[b] | Unmodified | 1 |
| 1500.766 | 25327 | 1500.767 | T81-K96+Na | Unmodified | 0 |
| 1501.769 | 16479 | 1501.770 | T81-K96+Na[b] | Unmodified | 0 |
| 1590.944 | 146309 | | | DMSO | |
| 1591.944 | 129634 | | | DMSO | |
| 1592.945 | 59199 | | | DMSO | |

[a]All m/z calculated for $[M+H]^+$ species unless otherwise indicated.

[b]Minor isotopomer. Isotopomer m/z determined using Protein Prospector.

[c]DMSO indicates that although the peak was identified it is also observed in DMSO control.

**Table S7 cont'd.** Fibril crosslinking with 100 μM EX-6-CLX

| Obs m/z | Intensity (Counts) | Calc m/z | Tryptic Peptide[a] | Notes[c] | Difference (ppm) |
|---|---|---|---|---|---|
| 1593.953 | 2822 | | | DMSO | |
| 1606.924 | 23077 | 1606.880 | T81-K97 | Unmodified | -28 |
| 1607.924 | 16553 | 1607.883 | T81-K97[b] | Unmodified | -26 |
| 1715.843 | 71953 | 1715.844 | E46-K58 | Modified | 0 |
| 1716.846 | 74342 | 1716.847 | E46-K58[b] | Modified | 0 |
| 1717.852 | 33744 | 1717.849 | E46-K58[b] | Modified | -1 |
| 1737.827 | 21631 | 1737.826 | E46-K58+Na | Modified | -1 |
| 1738.827 | 21382 | 1738.829 | E46-K58+Na[b] | Modified | 1 |
| 1884.838 | 173105 | | | DMSO | |
| 1885.839 | 178951 | | | DMSO | |
| 1886.839 | 94405 | | | DMSO | |
| 1887.841 | 32571 | | | DMSO | |
| 1928.046 | 312683 | 1928.045 | E61-K80 | Unmodified | -1 |
| 1929.046 | 328143 | 1929.048 | E61-K80[b] | Unmodified | 1 |
| 1930.045 | 175610 | 1930.050 | E61-K80[b] | Unmodified | 3 |
| 1931.046 | 64072 | 1931.053 | E61-K80[b] | Unmodified | 4 |
| 1932.052 | 7880 | 1932.056 | E61-K80[b] | Unmodified | 2 |
| 1950.018 | 48172 | 1950.027 | E61-K80+Na | Unmodified | 4 |
| 1951.024 | 46934 | 1951.030 | E61-K80+Na[b] | Unmodified | 3 |
| 1952.024 | 23017 | 1952.032 | E61-K80+Na[b] | Unmodified | 4 |
| 2196.062 | 86709 | 2196.111 | E44-K60+Na | Modified | 22 |
| 2197.062 | 102947 | 2197.114 | E44-K60+Na[b] | Modified | 24 |
| 2198.065 | 60902 | 2198.117 | E44-K60+Na[b] | Modified | 24 |
| 2199.067 | 19810 | 2199.119 | E44-K60+Na[b] | Modified | 24 |

[a]All m/z calculated for $[M+H]^+$ species unless otherwise indicated.

[b]Minor isotopomer. Isotopomer m/z determined using Protein Prospector.

[c]DMSO indicates that although the peak was identified it is also observed in DMSO control.

### Radioligand Binding

*Radioligand precursor synthesis*

Chemicals were obtained from commercial sources and used without further purification. Solvents were purchased from commercial sources and used as received unless stated otherwise. Reactions were performed at room temperature unless stated otherwise. Reactions were monitored by thin layer chromatography (TLC) on pre-coated silica 60 F254 aluminum plates (MilliporeSigma, Burlington, MA, USA), and spots were visualized by ultraviolet (UV) light. Evaporation of solvents was performed under reduced pressure at 40 °C using a rotary evaporator. Flash column chromatography was performed on a Biotage® (Charlotte, NC, USA) Isolera One system equipped with Biotage® SNAP KP-Sil cartridges. Nuclear magnetic resonance (NMR) spectroscopy was performed on a Bruker (Billerica, MA, USA) Avance Neo 600 (600 MHz for $^{1}H$ and 150 MHz for $^{13}C$) with chemical shifts (δ) reported in parts per million (ppm) relative to the solvent ($CDCl_3$, $^{1}H$ 7.26 ppm, $^{13}C$ 77.16 ppm; dimethyl sulfoxide (DMSO)-$d_6$, $^{1}H$ 2.50 ppm, $^{13}C$ 39.52 ppm). Low resolution liquid chromatography mass spectrometry (LCMS) was carried out using a Waters (Milford, MA, USA) SQD equipped with an Acquity UPLC instrument in positive-ion mode. High resolution mass spectrometry (HRMS) for small molecules was obtained on a Waters LCT Premier XE LC/MS system or Bruker Sci-Max. Python version 3.10.13 was used for data analysis scripts. ChatGPT drafted visualization scripts, and the scripts were checked and edited thoroughly for implementation details.

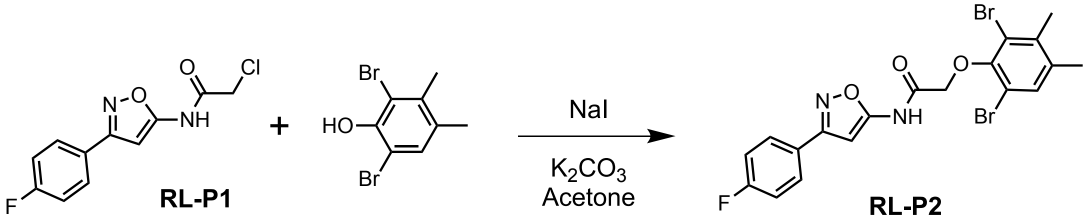


2-chloro-N-(3-(4-fluorophenyl)isoxazol-5-yl)acetamide (**RL-P1**) was synthesized as previously described (10). To a flame dried round bottom flask equipped with a stir bar, **RL-P1** (98.2 nmol, 1 equiv.), NaI (39.3 nmol, 0.5 equiv.), 2,6-dibromo-3,4-dimethylphenol (117.8 nmol, 1.5 equiv.), and $K_2CO_3$ (314.2 nmol, 4 equiv.) were added. The flask was set up with a reflux condenser and backfilled with argon (3x). Acetone (2 mL) was added to the flask and the reaction mixture was heated at reflux for four hours. The solvent was removed under reduced pressure. To the crude mixture, water (10 mL) was added and the aqueous layer was washed with EtOAc (10 mL x3). The combined organic layers were washed with brine (10 mL) and dried over $Na_2SO_4$. The solvent was removed under reduced pressure and purfied via flash chromotography utilizing EtoAC/Hexanes 10:90 (v/v) to obtain **RL-P2**.

**RL-P2** Characterization. Off-white solid; TLC (10:90 EtOAc:Hexanes v/v) $R_f$ 0.45; 95% yield. $^{1}H$ NMR (600 MHz, $CDCl_3$) δ 9.53 (s, 1H), 7.87 – 7.80 (m, 2H), 7.36 (s, 1H), 7.19 – 7.13 (m, 2H), 6.80 (s, 1H), 4.69 (s, 2H), 2.37 (s, 3H), 2.32 (s, 3H). $^{13}C$ NMR (150 MHz, $CDCl_3$) δ 164.79, 164.24, 163.13, 162.91, 159.83, 149.08, 137.38 (d, $J$ = 216.4 Hz), 133.02, 128.79 (d, $J$ = 8.3 Hz), 125.26 (d, $J$ = 3.0 Hz), 120.78, 116.04 (d, $J$ = 21.8 Hz), 113.41, 87.39, 70.42, 20.74, 19.89. HRMS (*m/z*): calculated for $C_{19}H_{16}Br_2FN_2O_3$ $[M+H]^+$ 496.9512, found 496.9524.

*Radioligand tritiation*

All solvents used were analytical grade and commercially available from Sigma-Aldrich and Fisher Scientific. Anhydrous solvents were routinely used for reactions. Reactions were typically run under an inert atmosphere of nitrogen or argon. Tritium gas was handled in a tritium gas manifold system (RC TRITEC AG, Teufen). Standard chemicals were bought from Sigma-Aldrich, Acros, Strem and Chemtronica AB (Stockholm). Mass spectra were recorded on a Waters Acquity UPLC/MS consisting of a Waters Binary Solvent Manager, Sample Manager, Column Manager, PDA Detector and a QDa Mass Detector in electrospray mode. An Acquity UPLC HSS (C18, 2.1 × 50 mm, 1.7 µm); the column temperature was set to 50 °C and the flow rate to 0.4 mL/min. A linear gradient was applied, starting at 95 % 10 mM ammonium hydrogencarbonate and ending at 97 % acetonitrile, in 4 min. HPLC analyses were performed on an Agilent 1100, HPLC system with a binary pump, auto-injector, DAD and column oven, coupled in series with a Packard Radiomatic Flow Scintillator 625TR, equipped with a liquid scintillation cell with a volume of 100 µL using UltimaFlow scintillation cocktail. An Atlantis T3 (C18, 4.6 × 100 mm, 3.5 µm); the column temperature was set to 40 °C and the flow rate to 0.8 mL/min. A linear gradient was applied, starting at 95% trifluoroacetic acid (0.1 %) and ending at 95 % acetonitrile, in 13 min. Preparative chromatography was run on a Gilson 305 with a Gilson UV/VIS-151 and a RAYTEST Ramona (Raytest Nordic AB, Höllviken, Sweden)equipped with Kromasil column (C18, 5 mm, 10 × 250 mm, C8, 7 mm, 10 × 250 mm or C8, 7 mm, 50 × 250 mm) using acetonitrile/50mM ammonium acetate in MilliQ Water from Pretech Instruments, (Sollentuna, Sweden). Liquid scintillation analysis was performed on a PACKARD TRI-CARB 2900TR from Chemical Instruments AB, CIAB, (LIDINGÖ, Sweden). Thin layer chromatography (TLC) was performed on Merck TLC-plates (Silica gel 60 F254) and UV- light (254 nm) visualized the spots. Flash column chromatography was performed on Silica gel 60.

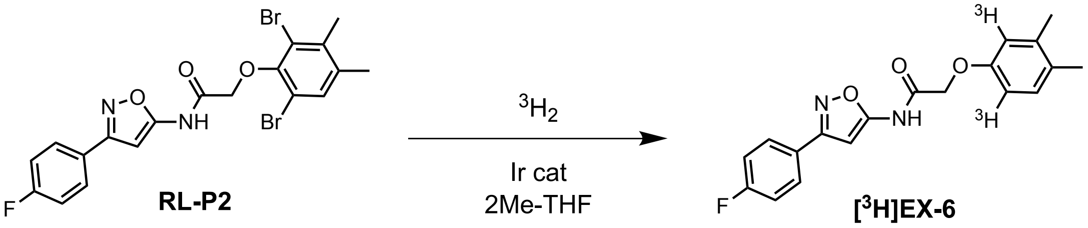


To a mixture of **RL-P2** (1.48 mg) and [(1,2,5,6-η)-1,5-cyclooctadiene][1,3-dihydro-1,3-bis(2,4,6-trimethylphenyl)-2H-imidazol-2-ylidene](dimethylphenylphosphine)-Iridium(**Ir cat**, 1.11 mg) 2-methyltetrahydrofuran (2-MeTHF, 400 µL) was added and flushed with tritium (628 mbar, 11 Ci). The mixture was stirred at ambient temperature overnight. The mixture was then filtered through a plug of silica (5 mm). Further elution was made with acetonitrile (2 × 2 mL) followed by evaporation of the solvents. The residue was dissolved in eluent (1.5 + 1.5 mL). Purification was performed by multiple injections on preparative reversed phase HPLC (KROMASIL C18, 7 µm, 250 × 10 mm using 75 % acetonitrile in 0.1 % TFA, 2.0 mL/min, UV 254 nm). Fractions with product were pooled and water (18 mL) was added. The solution was passed through a Sep-Pak (Sep-Pak® Plus Short tC18 Cartridge, washed with ethanol and water). The Sep-Pak was washed again with water (10 mL). The product (110 mCi) was extracted from Sep-Pak with ethanol. An aliquot (20.6 mCi) was diluted to a concentration of 1.0 mCi/mL; Molar activity: 56 Ci/mmol. When **[$^3$H]EX-6** is stored at -20 °C in its original solvent, the rate of decomposition is initially less than 2 % for 5 months.

**[$^3$H]EX-6** Characterization. Radiochemical purity (Radio-HPLC): >99 %; Chromatographic purity (220/254/280 nm): >99 %; Radiochemical concentration: 1.0 mCi/mL (38 MBq/mL); Molar activity: 56 Ci/mmol (2.09 TBq/mmol); LC-MS ID: Conforms with structure and EX-6 reference. **[$^3$H]EX-6** characterization data are shown in **Figure S19**.

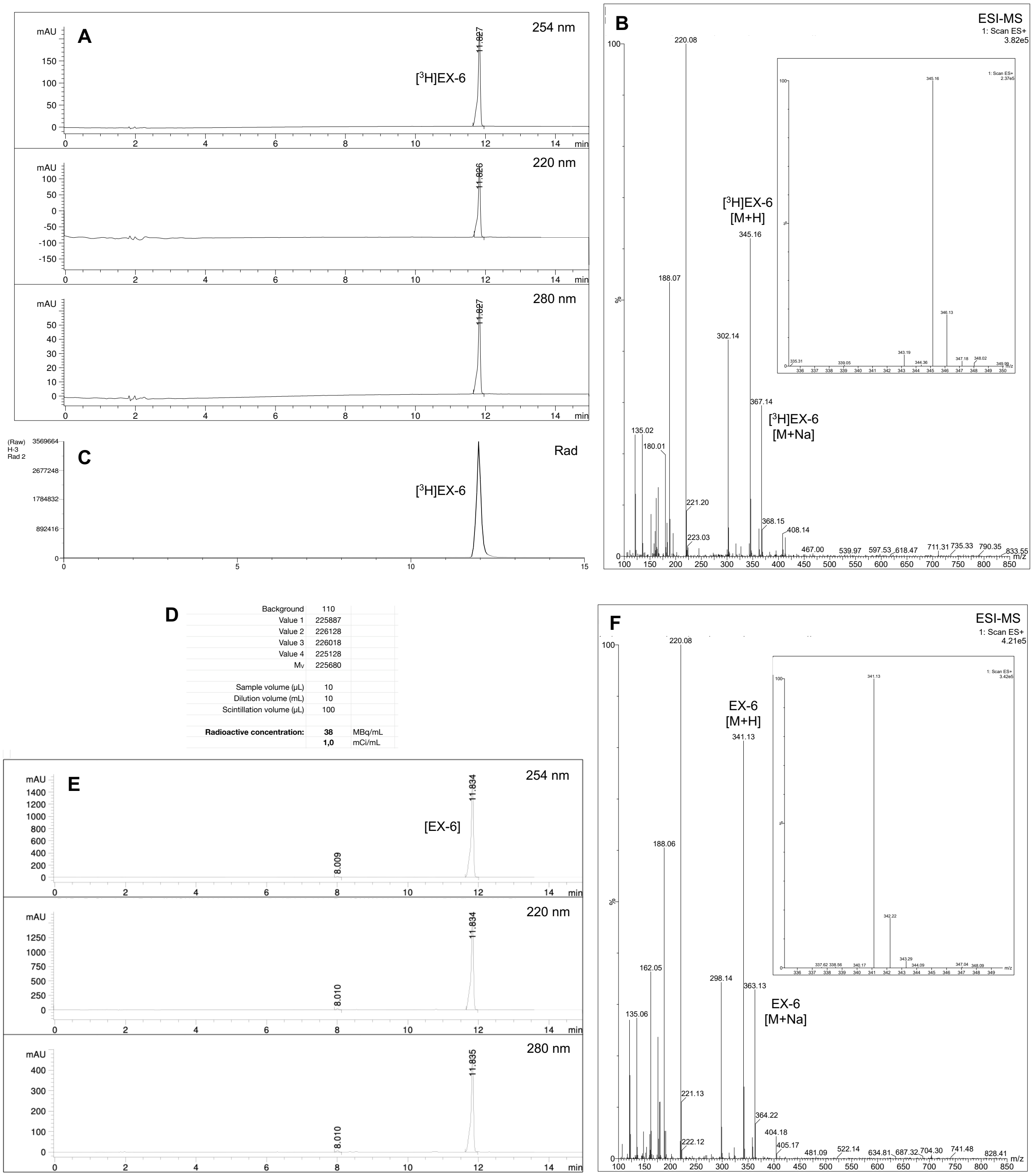


**Figure S17** Radioligand Characterization. A. UV traces from HPLC analysis of [$^3$H]EX-6. B. ESI positive ion MS data for [$^3$H]EX-6. C. Radioactivity trace from HPLC analysis of [$^3$H]EX-6. D. Radioactivity analysis of [$^3$H]EX-6. E. UV traces from HPLC analysis of EX-6. F. ESI positive ion MS data for EX-6.

*Autopsy material*
Formalin-fixed paraffin-embedded (FFPE) human brain tissue from Parkinson's disease (PD), dementia with Lewy bodies (DLB), multiple system atrophy (MSA), Parkinsonian-type (MSA-P) or cerebellar-type (MSA-C), Alzheimer's disease (AD), and control cases were acquired from the Center for Neurodegenerative Disease Research (CNDR) at the University of Pennsylvania. Refer to (**Table S9**) for the clinical demographic data of the cases used in this study. PD and MSA-P tissues for homogenate binding assays were frozen and acquired from the Banner Sun Health Research Institute (**Table S10**).

*Preparation of human brain tissue for in vitro binding studies*
As described in Bagchi et al. to prepare insoluble fraction from PD and MSA-P patients (11), fresh frozen tissue blocks were sequentially homogenized in four buffers (3 mL/g wet weight of tissue) with glass Dounce tissue grinders (Kimble): 1) High salt (HS) buffer: 50 mM Tris-HCl pH 7.5, 750 mM NaCl, 5 mM EDTA; 2) HS buffer with 1%Triton X-100; 3) HS buffer with 1%Triton X-100 and 1 M sucrose; and 4) phosphate buffered saline (PBS). Homogenates were centrifuged at 100,000 x g after each homogenization step, and the pellet was resuspended and homogenized in the next buffer in the sequence.

*Saturation binding assays*
Saturation binding assays were performed in MSA and PD brain homogenates (0.1 mg/mL tissue) using a [$^3$H]EX-6 concentration range of 1 nM to 150 nM, with incubation for 90 min at room temperature (RT) in PBS + 20% EtOH. Non-specific binding was determined using 10 μM of unlabeled EX-6. Bound and free radioligand is harvested with the Unifilter-96 harvesting system. (Perkin Elmer), followed by two washes with 250 μL ice-cold buffer containing PBS + 20 % EtOH. Filters containing bound ligands are added with 50 μL scintillation cocktail (MicroScint-20, PerkinElmer) and counted on the Microbeta system (PerkinElmer). The saturation data was fitted and analyzed using the non-linear regression function of GraphPad Prism 10 software to calculate the dissociation constant ($K_D$) and maximum number of binding sites ($B_{max}$). Scatchard plots were prepared with GraphPad Prism 10 software to display the saturation binding data.

*In vitro autoradiography and immunohistochemistry*
*In vitro* autoradiography was performed using 6-μm-thick FFPE sections derived from PD, DLB, AD, and elderly control brains. Brain sections underwent deparaffinization and an antigen retrieval step using 0.5 M citrate buffer (pH 6.0) for 1 hour at 70°C in a preheated water bath. All slides were subsequently equilibrated for 30 minutes in 1x PBS (Dulbecco's phosphate-buffered saline) and then incubated for 90 minutes at ambient temperature with 4.5 nM [$^3$H]EX-6. The sections were rinsed three times in cold buffer 1× PBS + 20% EtOH for 5 min, followed by a quick dip in cold distilled water. Non-specific binding was determined using 10 μM of unlabeled EX-6. Slides were then allowed to air-dry before being exposed and scanned in a real-time autoradiography system (BeaQuant instrument, ai4R) for 24 h. ROI delineation and quantification of signal were performed by using the image analysis software Beamage (ai4R). Specific binding was determined by subtracting the non-specific signal from the total signal and expressed as counts/min/mm$^2$. Immunohistochemistry was performed on deparaffinized sections adjacent to those used for autoradiography. Antigen retrieval for αSyn was performed by incubating the sections in 98% formic acid for 30 minutes and then steaming them in deionized water for 30 minutes, whereas Aβ sections were incubated in citrate buffer (0.5 M, pH 6.0) for 1 hour at 70 °C in a preheated water bath. Afterwards, sections were permeabilized with 0.1% Triton X-100 for 10 minutes, followed by three 5-minute washes in PBS-Tween 20 (PBST) buffer. Sections underwent hydrogen peroxide blocking for 15 min and PBS + 10% goat serum + 1% BSA + 0.1% Tween 20 blocking for 1 h at ambient temperature. Sections were immunostained using anti-Alpha-synuclein (pS129) antibody P-syn/81A and anti-beta Amyloid antibody [mOC23] (**Table S11**) used at 1:500 dilution overnight at 4 °C. After a series of thorough washes with PBST buffer, the slides were incubated with the secondary antibody Goat Anti-Mouse IgG H&L (HRP) and Goat anti-rabbit IgG H&L (HRP) (**Table S12**) at 1:10,000 for 1 h at ambient temperature. The sections were rewashed with PBST buffer three times for 5 minutes each, treated with DAB substrate for 10 minutes, and then counterstained with Meyer's hematoxylin dye. Subsequently, the sections were dehydrated sequentially into 50, 75, 95, and 100% ethanol and xylene baths for 5 min each, then mounted with Limonene mounting media and cover slipped for microscopy. Images were captured with a Leica Aperio slide scanner (RRID:SCR_022420) at 10x magnification. Tissue autoradiography and immunostaining data for PD, DLB,

AD, MSA-C, and control brain samples are shown in **Figure S20** and **Figure S21**. Quantified binding from all tissue samples is shown in **Figure S22**.

**Table S9** Selected demographics of cases used for autoradiography and immunohistochemistry.

| Case patient (P)/(C) | Gender (M/F) | Age of onset (Years) | Age at death (Years) | Tau | Aβ | αSyn | TDP-43 | Region used | Diagnosis |
|---|---|---|---|---|---|---|---|---|---|
| P | M | 80 | 85 | 0.5+ | 0 | 2+ | 0 | Cingulate cortex | PD |
| P | F | 55 | 62 | 0 | 0 | 3+ | 0 | Dentate nucleus | MSA-C |
| P | M | 51 | 59 | 0 | 0 | 3+ | 0 | Cingulate cortex | MSA-P |
| P | M | 65 | 83 | 0 | 3+ | 1+ | 0 | Cingulate cortex | DLB |
| P | M | 60 | 67 | 1+ | 3+ | 2+ | 0 | Cingulate cortex | DLB |
| P | M | 70 | 73 | 1+ | 3+ | 3+ | 0 | Cingulate cortex | DLB |
| P | F | 67 | 74 | 3+ | 3+ | 0 | 0 | Frontal | AD |
| C | M | | 66 | 0 | 0 | 0 | 0 | Cingulate gyrus | CT |

**Table S10** Demographic data of cases from Banner Sun Health Research Institute selected for brain homogenates.

| Diagnosis | Sex(M/F) | Age (years) | PMT | LB stage |
|---|---|---|---|---|
| PD | M | 81 | 2.5 | III. Brainstem/Limbic |
| PDD | M | 70 | 1.83 | III. Brainstem/Limbic |
| PDD | M | 75 | 2.25 | IV. Neocortical |
| PD | M | 75 | 2.33 | IV. Neocortical |
| MSA-P | F | 57 | 32 | n/a. |
| MSA-P | M | 58 | 12 | n/a |

**Table S11** Primary antibody selected for immunohistochemistry.

| Primary Antibodies | Source/Cat No | Host and clonality | Antibody dilution used | Incubation time |
|---|---|---|---|---|
| Anti-alpha-synuclein (phospho S129) antibody P-syn/81A | Abcam/ab184674) | Mouse monoclonal | 1:500 | Overnight at 4 ºC |
| Anti-beta Amyloid antibody [mOC23] | Abcam/ab205340 | Rabbit monoclonal | 1:500 | Overnight at 4 ºC |

**Table S12** Secondary antibody selected for immunohistochemistry.

| Secondary Antibody | Source/Cat No | Host and clonality | Antibody dilution used | Incubation time |
|---|---|---|---|---|
| Goat Anti-Mouse IgG H&L | Abcam/ab205719 | Goat polyclonal | 1:10000 | 1 h at ambient temperature |
| Goat Anti-Rabbit IgG H&L | Ab97051 | Goat polyclonal | 1:10000 | 1 h at ambient temperature |

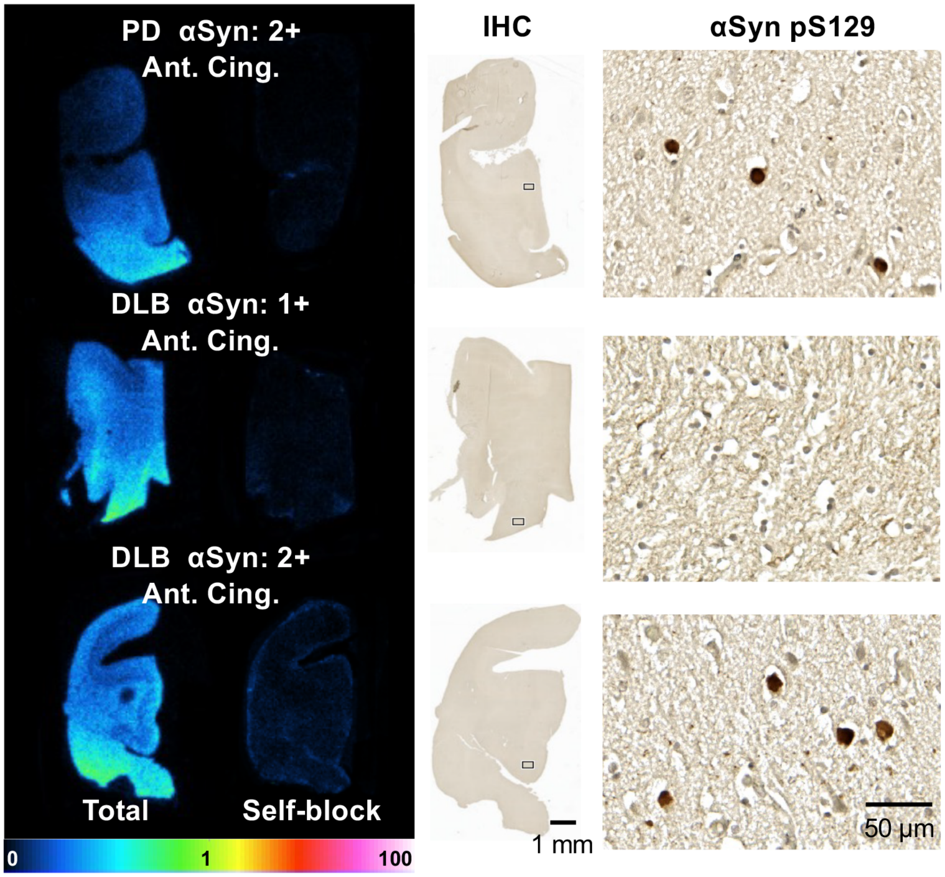


**Figure S18** Radioligand Binding Data 1. Tissue autoradiography studies with [$^{3}$H]EX-6 demonstrating selectivity for binding to PD and DLB cases with IHC staining for αSyn pS129.

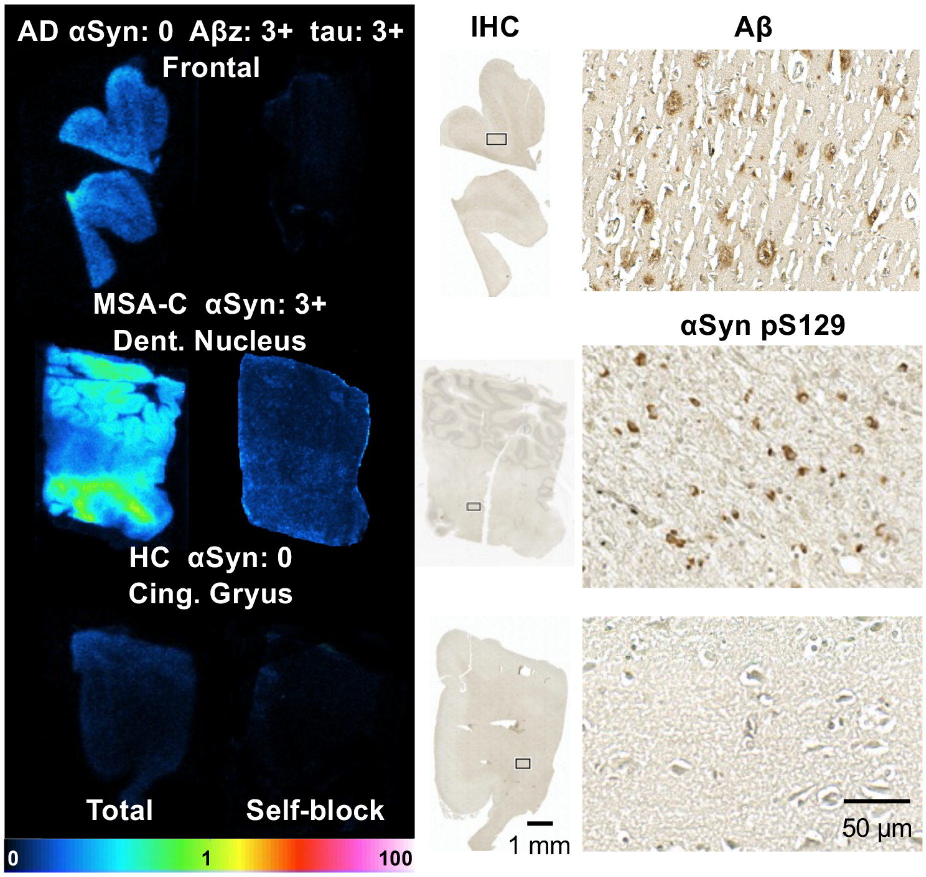


**Figure S19** Radioligand Binding Data 2. Tissue autoradiography studies with [$^3$H]EX-6 demonstrating selectivity for binding to AD, MSA-C, and HC cases with IHC staining for Aβ or αSyn pS129.

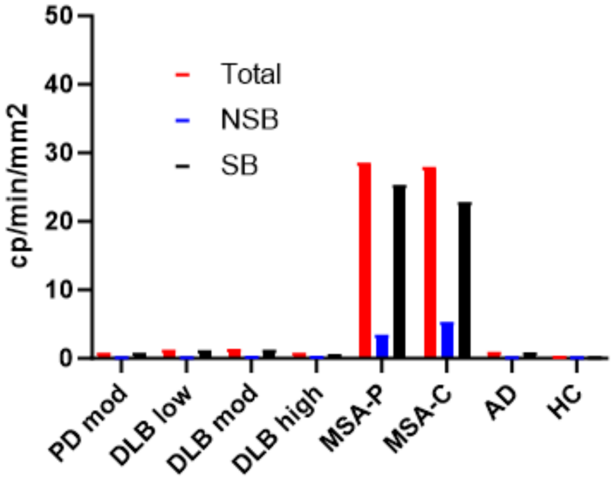


**Figure S20** Radioligand Binding Summary. Total binding, non-specific binding (NSB), and specific binding (SB) from tissue autoradiography studies with [$^3$H]EX-6 demonstrating selectivity for binding to AD, MSA-C, and HC cases.

### References


1. M. D. Tuttle *et al.*, Solid-state NMR structure of a pathogenic fibril of full-length human alpha-synuclein. *Nat Struct Mol Biol* **23**, 409-415 (2016).
2. C.-J. Hsieh *et al.*, Alpha Synuclein Fibrils Contain Multiple Binding Sites for Small Molecules. *ACS Chemical Neuroscience* **9**, 2521-2527 (2018).
3. L. C. Speight *et al.*, Efficient Synthesis and In Vivo Incorporation of Acridon-2-ylalanine, a Fluorescent Amino Acid for Lifetime and Förster Resonance Energy Transfer/Luminescence Resonance Energy Transfer Studies. *Journal of the American Chemical Society* **135**, 18806-18814 (2013).
4. J. G. Marmorstein *et al.*, Improved Large-Scale Synthesis of Acridonylalanine for Diverse Peptide and Protein Applications. *Bioconjugate Chemistry* **35**, 1913-1922 (2024).
5. J. J. Ferrie *et al.*, Using a FRET Library with Multiple Probe Pairs To Drive Monte Carlo Simulations of α-Synuclein. *Biophysical Journal* **114**, 53-64 (2018).
6. M. B. Cory *et al.*, FRETing about the details: Case studies in the use of a genetically encoded fluorescent amino acid for distance-dependent energy transfer. *Protein Science* **32**, e4633 (2023).
7. A. V. West *et al.*, Labeling Preferences of Diazirines with Protein Biomolecules. *Journal of the American Chemical Society* **143**, 6691-6700 (2021).
8. Y. Jiang *et al.*, Dissecting diazirine photo-reaction mechanism for protein residue-specific cross-linking and distance mapping. *Nature Communications* **15**, 6060 (2024).
9. UCSF (Protein Prospector.
10. M. G. Lougee *et al.*, Harnessing the intrinsic photochemistry of isoxazoles for the development of chemoproteomic crosslinking methods. *Chemical Communications* **58**, 9116-9119 (2022).
11. D. P. Bagchi *et al.*, Binding of the Radioligand SIL23 to α-Synuclein Fibrils in Parkinson Disease Brain Tissue Establishes Feasibility and Screening Approaches for Developing a Parkinson Disease Imaging Agent. *PLOS ONE* **8**, e55031 (2013).
